# Auditory Activity is Diverse and Widespread Throughout the Central Brain of *Drosophila*

**DOI:** 10.1101/709519

**Authors:** Diego A Pacheco, Stephan Y Thiberge, Eftychios Pnevmatikakis, Mala Murthy

## Abstract

Sensory pathways are typically studied by starting at receptor neurons and following postsynaptic neurons into the brain. However, this leads to a bias in analysis of activity towards the earliest layers of processing. Here, we present new methods for volumetric neural imaging with precise across-brain registration, to characterize auditory activity throughout the entire central brain of *Drosophila* and make comparisons across trials, individuals, and sexes. We discover that auditory activity is present in most central brain regions and in neurons known to carry responses to other modalities. Auditory responses are temporally diverse, but the majority of activity, regardless of brain region, is tuned to aspects of courtship song. We find that auditory responses are stereotyped across trials and animals in early mechanosensory regions, becoming more variable at higher layers of the putative pathway, and that this variability is largely independent of spontaneous movements. This study highlights the power of using an unbiased, brain-wide approach for mapping the functional organization of sensory activity.

## INTRODUCTION

A central problem in neuroscience is determining how information from the outside world is represented by the brain. Solving this problem requires knowing which brain areas and neurons encode particular aspects of the sensory world, how representations are transformed from one layer to the next, and the degree to which these representations vary across presentations and individuals. Most sensory pathways have been successfully studied by starting at the periphery – the receptor neurons - and then characterizing responses along postsynaptic partners in the brain. We therefore know most about the earliest stages of sensory processing in most model organisms. However, even for relatively simple brains, there exists a significant amount of cross-talk between brain regions ^1, 2^. Thus, neurons and brain areas may be more multi-modal than appreciated. Here, we present methods to systematically characterize sensory responses throughout the *Drosophila* brain, for which abundant neural circuit tools facilitate moving from activity maps to genetic reagents that target identifiable neurons ^3^.

We focus on auditory coding in the *Drosophila* brain because, despite its relevance to courtship behavior, surprisingly little is known regarding how auditory information is processed downstream of primary and second order mechanosensory neurons. In addition, in contrast with olfaction or vision, for which the range of odors or visual patterns flies experience in naturalistic conditions is large, courtship song is the main auditory cue to which flies respond, and it is comprised of a narrow range of species-specific features that can be easily probed in experiments ^4^. *Drosophila* courtship behavior unfolds over many minutes, and females must extract features from male songs to inform mating decisions ^5, 6^ while males listen to songs to inform their courtship decisions in group settings ^7^. Fly song has only three modes, two types of brief sound impulses or ‘pulse song’ and one hum-like ‘sine song’^8^, and males both alternate between these modes to form song bouts and modulate song intensity based on sensory feedback ^9, 10^.

Flies detect sound using a feathery appendage of the antenna, the arista, which vibrates in response to near field sounds ^11^. Antennal displacements activate mechanosensory Johnston’s Organ neurons (JONs) housed within the antenna. Three major populations of JONs (A, B, and D) respond to vibratory stimuli at frequencies found in natural courtship song ^12, 13^. These neurons project to distinct areas of the antennal mechanosensory and motor center (AMMC) in the central brain ^14^. Recent studies suggest that the auditory pathway continues from the AMMC, to the wedge (WED), then to the ventrolateral protocerebrum (VLP), and to the lateral protocerebral complex (LPC) ^4, 15–19^. However, our knowledge of the fly auditory pathway remains incomplete and the functional organization of regions downstream of the AMMC and WED are largely unexplored. Moreover, nearly all studies of auditory coding in *Drosophila* have been performed on female brains, even though both males and females process courtship song information ^4^.

To address these issues, we developed methods to investigate the representation of behaviorally relevant auditory signals throughout the central brain of *Drosophila*, and to make comparisons across animals. We image activity volumetrically combined with fully automated segmentation of regions of interest ^20^. We use GCaMP6s and two-photon microscopy to sequentially target the entirety of the *Drosophila* central brain *in vivo*. In contrast with recent brain-wide imaging studies in *Drosophila* ^1, 21^, we have traded off temporal speed for enhanced spatial resolution. Imaging at high spatial resolution facilitates automated region of interest (ROI) segmentation, each ROI covering sub-neuropil structures, including cell bodies and neurites. ROIs were accurately registered into an *in vivo* template brain, to compare activity across trials, individuals, and sexes, and to build comprehensive maps of auditory activity throughout the central brain.

Our results reveal that the representation of auditory signals is broadly distributed throughout 33 out of 36 major brain regions, including in regions known to process other sensory modalities, such as all levels of the olfactory pathway, or to drive various motor behaviors, such as the central complex. The representation of auditory stimuli is diverse across brain regions, but focused on conspecific features of courtship song. Auditory activity is more stereotyped (across trials and individuals) at early stages of the putative mechanosensory pathway, becoming more variable and more selective for particular aspects of the courtship song at higher stages. This variability cannot be explained by simultaneous measurements of fly behavior. On the other hand, auditory maps are largely similar between male and female brains, despite extensive sexual dimorphisms in neuronal number and morphology. These findings provide the first brain-wide description of sensory processing and feature tuning in *Drosophila*.

## RESULTS

### A new pipeline for mapping sensory activity throughout the central brain of *Drosophila*

We developed a pipeline to volumetrically image sensory responses (via GCaMP6s ^22^ expressed pan-neuronally), and then precisely register sequentially imaged volumes into an *in vivo* template [Figure 1A; Figure S1A]. We presented three distinct auditory stimuli, sinusoids, pulse trains, and broadband noise (see Methods). The first two stimuli represent the two major modes of *Drosophila melanogaster* courtship song - all three stimuli should drive auditory neurons, but not neurons sensitive to slow vibrations, such as those induced by wind or gravity ^12, 13^. For each fly, we imaged 14-17 subvolumes at 1Hz (voxel size 1.2×1.4×2 μm^3^) until covering the whole extent of the Z axis per fly (posterior to anterior), and ¼ of the central brain in X-Y - what we refer to hereafter as a “volume”. These volumes were time-series motion corrected and stitched along the posterior-anterior axis using a secondary signal ^23^ [Figure 1A-i]. We imaged volumes from either dorsal or ventral quadrants, providing full hemisphere coverage by imaging only two flies [Figure 1B]. We mirrored each volume such that all dorsal or ventral volumes imaged could be compared in the same reference brain coordinates (see below) - this is why all activity maps appear on ½ of the brain (e.g. [Figure S1B]).

**Figure 1.**
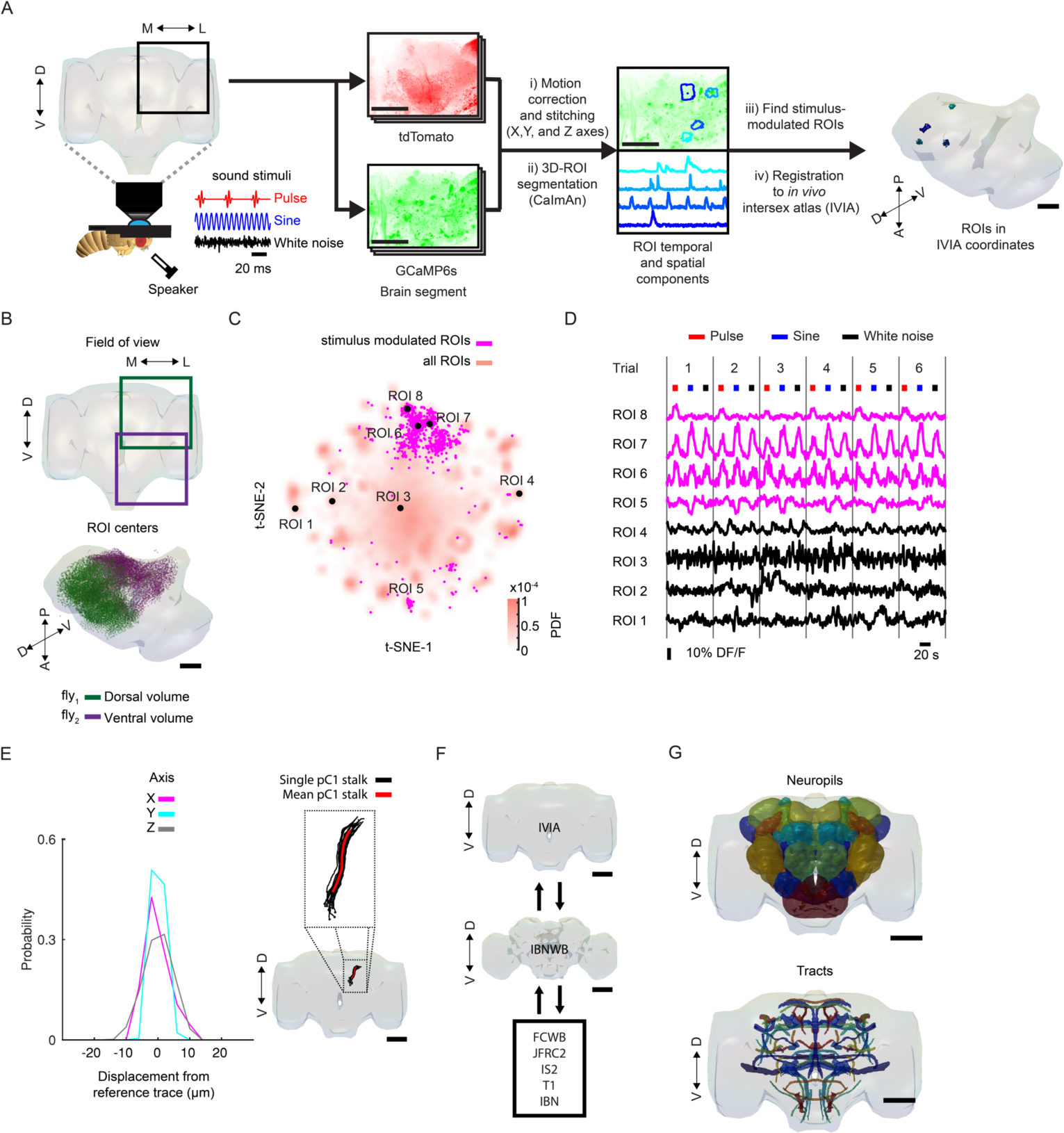
A new pipeline for mapping auditory activity throughout the central brain. (A) Overview of data collection and processing pipeline (for more details see Figure S1A): (i) tdTomato signal is used for motion correction of volumetric time-series and to stitch serially imaged overlapping brain segments in X, Y, and Z axes. (ii) 3D-ROI segmentation (via CaImAn) is performed on GCaMP6 signal. (iii) auditory ROIs are selected. (iv) ROIs are mapped to the *in vivo* intersex atlas (IVIA) space. (B) Example of 11,225 segmented ROIs combined from two flies (dorsal (green) and ventral (purple) volumes from each fly). ROI centers (dots, below) span the entire anterior-posterior and dorsal-ventral axis. These ROIs have not yet been sorted for those that are auditory (see Figure S1D-E). (C) 2D t-SNE embedding of activity from all ROIs in panel (B). ROIs modulated by auditory stimuli (1,118 out of 11,225 ROIs) are shown in magenta. (D) DF/F from ROIs indicated in (C). Magenta traces correspond to stimulus-modulated ROIs, while black traces are non-modulated ROIs. Time of individual auditory stimulus delivery is indicated (pulse, sine, or white noise). (E) (left) IVIA registration accuracy - per-axis jitter (X, Y, and Z) between traced pC1 stalk across flies relative to mean pC1 stalk (see Figure S2 for more details). (right) 3D-rendering of traced stalks of Dsx+ pC1 neurons (black traces, n = 20 brains from Dsx-GAL4/UAS-GCaMP6s flies) and mean pC1 stalk (red trace). (F) Schematic of bridging registration between IVIA and IBNWB atlas (*Ito et al. 2014*). This bidirectional interface provides access to a network of brain atlases (IBN, FCWB, JFRC2, IS2, and T1) associated with different *Drosophila* neuroanatomy resources (Manton et al. 2014). See Figure S3 for more details. (G) Anatomical annotation of IVIA. Neuropil and neurite tract segmentations from IBN mapped to IVIA (for a full list of neuropil and neurite tract names see Table S1). Scale bars in all panels, 100 μm. See also Movies S1-4.

To automate the segmentation of regions of interest (ROIs) throughout the brain, we used a constrained non-negative matrix factorization algorithm optimized for 3D time series data ^20, 24^ (see Methods) [Figure 1A-ii]. This method groups contiguous pixels that are similar in their temporal responses; thus, each ROI is a functional unit. We remain agnostic as to the specific relationship between ROIs and neurons - a single ROI could represent activity from many neurons (e.g., a cluster of axon tracts with similar responses), or a single neuron can be segmented into multiple ROIs if different neuronal compartments differ in their activity. Importantly, this method enables effective segmentation of responses from neuropil (axons, dendrites, and processes), not only somas - this is critical for capturing sensory responses in *Drosophila*, for which somas may not reflect activity in processes ^25^. We typically extracted thousands of ROIs per volume, and collectively sampled activity from all central brain regions [Figures 1C and S1B-C]. These ROIs included both spontaneous and stimulus-driven activity [Figure 1C]; stimulus-driven ROIs were identified using quantitative criteria (see Methods) [Figures 1A-iii and S1D-E]. These criteria select ROIs with responses consistent across trials [Figure 1D] - but see below for analysis of trial-to-trial variation even given these criteria [Figures 5-6]. We found hundreds of auditory ROIs per fly ranging from 1.2% to 25.4% of all segmented ROIs per fly [Figure S1F].

To compare activity across individuals and sexes, we next registered functionally imaged brain segments into an *in vivo* intersex atlas using the computational morphometry toolkit (see Methods) [Figure 1A-iv; Figures S1A and S2A]. Registration of individual fly brains to our *in vivo* intersex atlas (IVIA) is at cellular accuracy (∼4.3 um, across all tracts), measured as the deviation from the mean trace of several identifiable neural tracts (e.g., pC1 neurons, Figure 1E, and pC2l, pC2m, pMP7 neurons, Figure S2B). Using this approach, we successfully registered volumes from 45 out of 48 flies (see examples in Figure S2C).

The IBNWB *Drosophila* fixed brain atlas ^26^ contains detailed brain neuropil and neurite tract segmentation information. By linking our *in vivo* intersex template (IVIA) to the IBNWB atlas, we could map auditory activity onto known brain neuropils and processes [Figures 1F-G; Supplemental Table S1], and thereby link to the whole network of *D. melanogaster* anatomical repositories ^3^. We built a bridge registration between IVIA and the IBNWB atlas using a point set registration algorithm ^27^ with high accuracy (deviation of ∼2.24 um for IVIA-to-IBNWB, and ∼2.8 um for IBNWB-to-IVIA) (see Methods, [Figures S3A-C]).

In summary, we have developed a novel open-source pipeline for monitoring, anatomically annotating, and directly comparing sensory-driven activity, at high spatial resolution, across trials, individuals, and sexes.

### Auditory activity is widespread across the central brain

We extracted 19,036 auditory ROIs from 33 female and male brains [Figures 2A and S4A]. Within a single fly, auditory ROIs were broadly distributed across the anterior-posterior and dorsal-ventral axes [Figure 2B]. Probability density maps across flies using the ROIs’ spatial components (see Methods) reveal across-individual consistency of this broad auditory-driven activity [Figure 2C] - the spatial map was also similar between males and females [Figure S4B]. We found that 77% of all segmented ROIs were assigned to an identifiable neuropil or tract [Figures 2D-F and S4C-D]. Surprisingly, 33 out of 36 central brain neuropils contained robust auditory activity (see Methods). In addition, testing other song-relevant stimuli (natural song and synthetic pulse train with pulses of higher frequency ^8^) confirmed that we are not missing auditory responses narrowly selective to additional frequencies and patterns not present in the synthetic sine and pulse song we used as stimuli [Figure S5].

**Figure 2.**
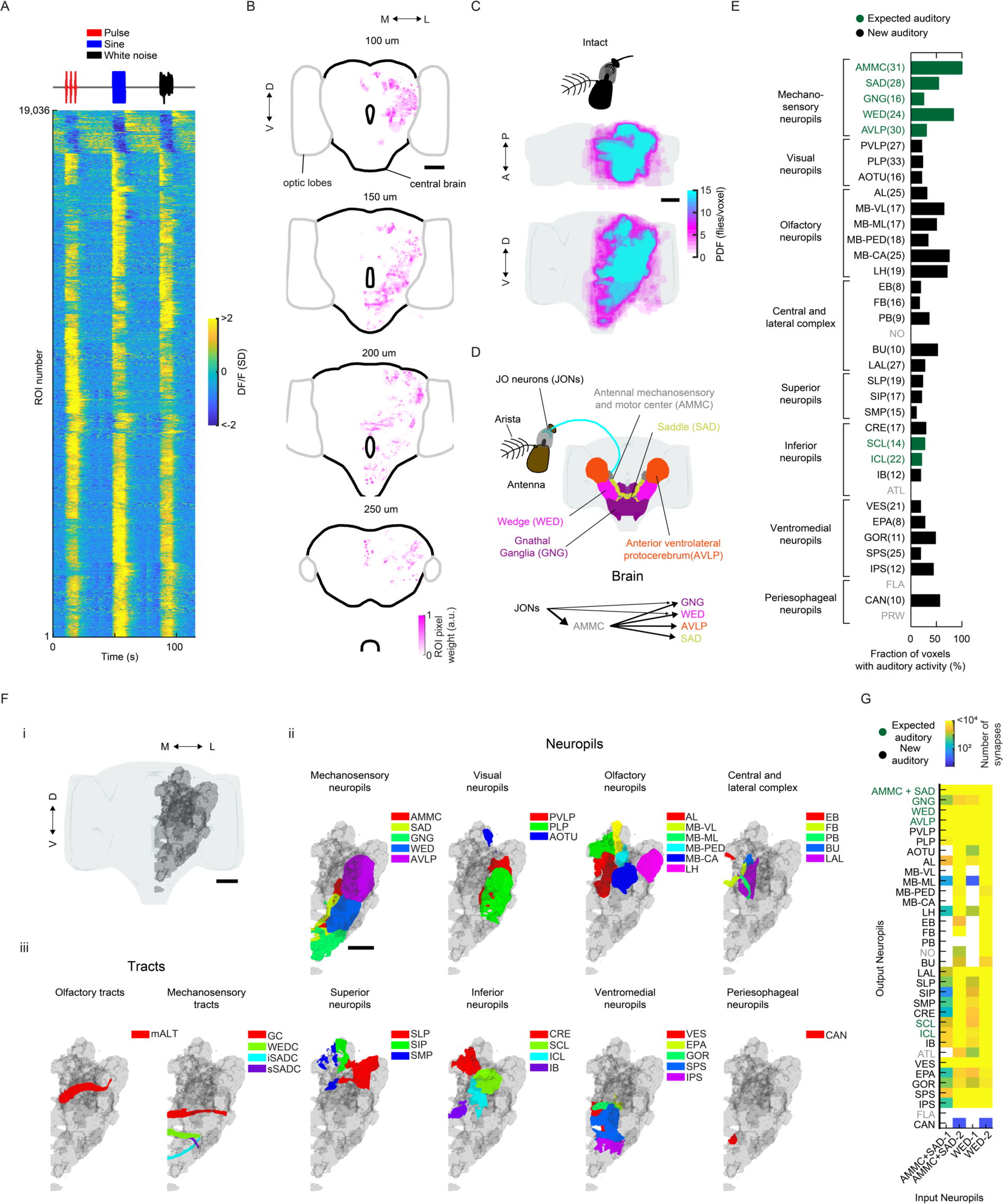
Auditory activity is widespread throughout the central brain of *Drosophila*. (A) Auditory ROI responses to pulse, sine, and white noise stimuli (n = 33 flies, 19,036 ROIs). Each row is the median across 6 trials - all responses are z-scored and therefore plotted as the change in the standard deviation (SD) of the DF/F signal. ROIs are sorted based on clustering of temporal profiles (see Figure 3A). (B) Spatial distribution of auditory ROIs (ROI pixel weights in magenta) combined from two flies (one ventral and one dorsal volume) (ROIs are in IVIA coordinates) - four different depths are shown (0 μm is the most anterior section of the brain and 300 um the most posterior). Grey contour depicts the optic lobes; black contour depicts the central brain. (C) Maximum projection (from two orthogonal views) of the density of auditory ROIs (n = 33 flies, 19,036 ROIs) - color scale is the number of flies with an auditory ROI per voxel. (D) Schematic of the canonical mechanosensory pathway in *D. melanogaster*. JONs in the antenna project to the AMMC, GNG and WED. AMMC neurons connect with the GNG, WED, SAD, and AVLP. (E) Fraction of voxels with auditory activity by central brain neuropil; percentages averaged across 33 flies (a minimum of 4 flies with auditory activity in a given neuropil was required for inclusion). The number of flies with auditory responses in each neuropil are indicated in parentheses. Neuropils with no auditory activity are indicated in gray font (also for panel G). (F) 3D-rendering of the volume that contains all auditory voxels across all flies (gray) and the overlap of this volume with brain neuropils (ii) and tracts (iii). Scale bars in all panels, 100 μm. (G) Using the ‘hemibrain’ connectomic dataset (*Scheffer et al. bioRxiv 2020*), heatmap of neuropils in which all identified AMMC/SAD or WED neurons (1079 and 3556, respectively) have direct synaptic connections (AMMC+SAD-1 or WED-1). Neuropils with connections to AMMC/SAD or WED neurons via one intermediate synaptic connection (AMMC+SAD-2 or WED-2). Neuropils with less than 5 synapses are shaded white. See also Figures S4, S5, S6, S7, and Movie S5.

We found auditory activity in all neuropils innervated by neurons previously described as auditory or mechanosensory ^4, 6, 15–17, 19, 28–30^, in the antennal mechanosensory and motor center (AMMC), saddle (SAD), wedge (WED), anterior ventrolateral protocerebrum (AVLP), gnathal ganglion (GNG), inferior (ICL), and superior clamp (SCL) - herein referred to as ‘expected auditory’ neuropils [Figures 2D-E]. Similarly, we detected auditory ROIs in many tracts or commissures (great commissure (GC), wedge commissure (WEDC), inferior (iSADC) and superior saddle commissure (sSADC)) connecting expected auditory neuropils [Figures 2F-iii and S4E-F].

Surprisingly, we found that a number of neuropils and tracts outside the expected auditory neuropils aforementioned contained a large percentage of auditory responses (fraction of voxels with auditory activity [Figure 2E]) - for example, neuropils of the olfactory pathway (antennal lobe (AL), the medial antennal lobe tract (mALT) containing olfactory projection neurons (PNs), the mushroom body (MB), including the peduncle, lobes, and calyx, and the lateral horn (LH)), parts of the visual pathway (posterior ventrolateral protocerebrum (PVLP) and posterior lateral protocerebrum (PLP)), and the central complex (ellipsoid body (EB), fan-shaped body (FB), protocerebral bridge (PB), bulb (BU), and lateral accessory lobe (LAL)) - such neuropils as herein referred to as ‘new auditory’ neuropils [Figures 2E and 2F]. We confirmed that activity within the olfactory pathway was present within the somas of individual antennal lobe projection neurons and mushroom body Kenyon cells [Figure S6]. Some of this new activity was in neuropils known to be downstream targets of ‘expected auditory’ neuropils but never previously studied functionally (e.g., the inferior posterior slope (IPS) and gorget (GOR) ^18^). These results demonstrate that auditory-evoked activity is widespread throughout the brain, extending beyond expected auditory neuropils, including in the olfactory, visual and pre-motor pathways.

### Widespread auditory activity originates in mechanosensory receptor neurons

To determine how auditory information makes its way to so many neuropils, we used the new ‘hemibrain’ connectomic resource ^31^. Mechanosensory receptor neurons (Johnston’s Organ neurons) project to the AMMC/SAD, and AMMC/SAD neurons connect with the WED ^14, 15, 18^. We determined the primary and secondary target neuropils for all AMMC/SAD and WED neurons identified in the hemibrain - this analysis revealed that neurons that originate in the AMMC/SAD or WED directly connect with neurons in 22 or 24 neuropils, and within one additional synapse, connect with neurons in 32 or 33 neuropils, respectively [Figure 2G].

To confirm that this activity was in fact driven by mechanosensory receptor neurons, we imaged flies carrying the *iav*^1^ mutation, rendering flies deaf ^32, 33^. This manipulation resulted in a 99% reduction in activity [Figures S7A-B]. Response magnitudes of the small number of auditory ROIs found in *iav*^1^ flies were significantly reduced compared with wild type flies [Figure S7C]. We thus conclude that the overwhelming majority of auditory activity we map with our stimuli are transduced through the mechanoreceptor neurons.

### Auditory activity throughout the central brain is characterized by diverse temporal responses

We next hierarchically clustered responses from auditory ROIs based on their temporal profiles (see Methods), and identified 18 distinct stimulus-locked response types [Figure 3A]. These same 18 response types are present in both male and female brains [Figure S8A], with only minor differences in central brain auditory representations between sexes [Figures S8B-C] – we therefore pooled male and female auditory ROIs for subsequent analyses. We measured the frequency of response types for each neuropil to evaluate differences in auditory representations by neuropil [Figure 3B]. Auditory activity is diverse throughout the central brain, consisting of both inhibitory (types 1-3) and excitatory (types 5-18) responses, along with diversity in response kinetics [Figures 3D-F]. Response types also differed in their selectivity for auditory stimuli [Figure 3G]. We interpret white noise preferring responses as a preference for frequencies outside of the range of the two fly courtship song stimuli tested (see also Figure 6).

**Figure 3.**
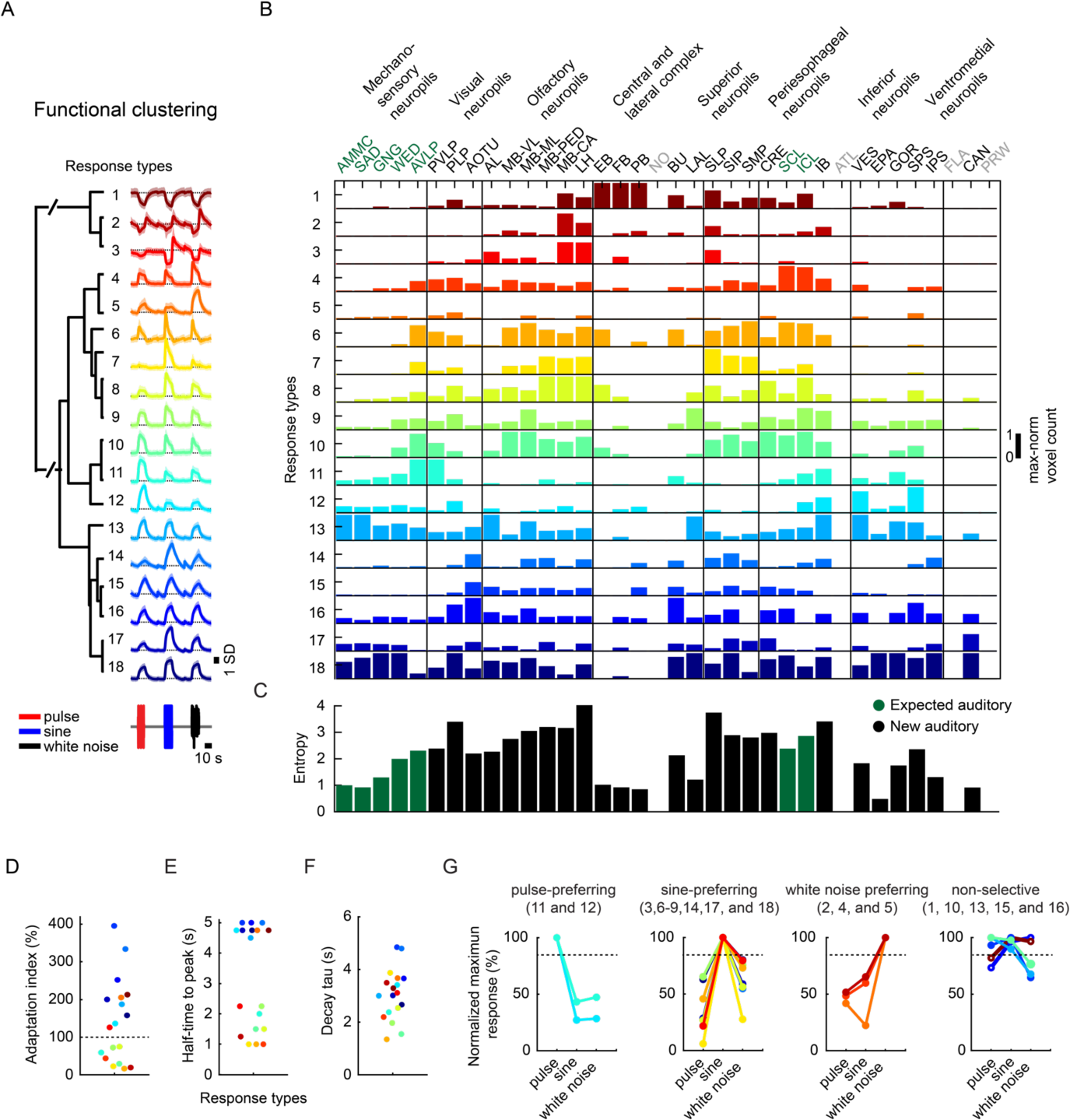
Brain-wide auditory activity is characterized by a diversity of temporal response profiles across neuropils. (A) Hierarchical clustering of auditory responses into 18 distinct response types (see Methods). Thick traces are the mean response (across ROIs) to pulse, sine, and white noise, and shading is the standard deviation (n = 33 flies, 19,036 ROIs). All responses are z-scored, and therefore plotted as the standard deviation (SD) of the responses over time, black dash line corresponds to mean baseline across stimuli for each response type. (B) Distribution of 18 response types across 36 central brain neuropils. Histogram of voxel count (how many voxels per neuropil with auditory activity of a particular response type) is max-normalized for each neuropil (normalized per column). Colorcode is the same as in (A) (n = 33 flies). Neuropils with no auditory activity are indicated in gray font. (C) Diversity (measured as the entropy across response type distributions shown in (B)) of auditory responses across central brain neuropils (see Methods). (D-F) Kinetics of all auditory responses by response type. Adaptation (D), Half-time to peak (E), and decay time tau (F). (G) Diversity in tuning for response types. i) pulse-preferring, ii) sine-preferring, iii) white noise preferring, and iv) non-selective response types (preference is defined by the stimulus that drives the maximum absolute response (at least 15% greater than the second highest response). Non-selective responses are divided into sine-and-pulse preferring (solid circles) and sine-and-white-noise preferring (open circles). See also Figures S8, S9, S10, and S11.

The diversity of response types found [Figure 3B] suggests that our method samples activity from many different neurons per neuropil. In addition, each response type had a distinct spatial distribution [Figures S9A-B]. We measured the entropy of response types per neuropil as a proxy for the diversity of auditory activity; responses were less diverse in the earliest mechanosensory areas and became more diverse at higher levels of the putative pathway [Figures 2D and 3C]. Response diversity was highest in new auditory areas, such as the posterior lateral protocerebrum (PLP), lateral horn (LH), and superior lateral protocerebrum (SLP). Clustering the data in Figure 3B revealed neuropils with highly similar response profiles, suggesting functional interconnectivity (e.g., AMMC/SAD or AVLP/PVLP or GNG/WED/GOR) [Figure S10].

Within the expected auditory neuropils [Figure 2D], we found that the AMMC contained predominantly non-selective (types 13 and 16) and sine-preferring (type 18) response types, but that brain regions downstream of the AMMC, showed changes in the distribution of response types, such as a decrease in non-selective response types and an increase in both white noise- and pulse-preferring response types [Figure 3B]. Prior work on individual neurons innervating the AVLP suggested this brain area mostly encodes pulse-like stimuli ^6, 15, 17^. We found, in contrast, strong representation of *both* sine- and pulse-preferring response types, with responses becoming more narrowly tuned in these regions (e.g., response types 7 and 11). In the lateral junction (SCL and ICL), as expected from innervations of pulse-tuned neurons^4^, there are a number of pulse-preferring response types (types 11 and 12). However, the predominant response types are sine-preferring.

In the new auditory neuropils, we found that the olfactory system was dominated by non-selective and sine-preferring response types, while pulse-preferring response types were mostly absent [Figures 3B, S6A-D]); in addition, response types became more diverse in higher-order olfactory regions [Figure 3C]. In the visual neuropils (PVLP, PLP, and AOTU), auditory activity was highly compartmentalized. For example, in the posterior ventrolateral protocerebrum (PVLP), activity was concentrated in the vicinity of ventro-medial optic glomeruli (LC4, LPLC1 and LPLC2), while in the PLP, auditory ROIs formed a single discrete compartment - minimally overlapping with optic glomeruli - that runs from the anterior WED-PLP boundary to a more posterior PLP-LH boundary [Figure S11]. In the central complex (EB, FB and PB) we found primarily non-selective and inhibitory response types (type 1), while the lateral complex (BU and LAL) contained both sine-tuned (types 18 and 6) and non-selective excitatory response types (types 16 and 13) [Figure 3B].

In summary, auditory activity across the central brain comprises at least 18 distinct response types, with both expected and new auditory neuropils showing a diversity of response types, and with responses becoming more stimulus-selective in deeper brain neuropils. Next, we investigate the tuning of auditory responses by presenting a larger battery of stimuli.

### Widespread auditory activity is centered on features of conspecific courtship songs

To assay tuning for courtship song features [Figure 4A], we focused on the neuropils with strongest auditory activity [Figure 2E; Figure 4B], and imaged subregions of each of these neuropils [Figure S12A], in order to sample ROIs with varying degrees of pulse or sine selectivity. Tuning curves were formed for each ROI and for each set of stimuli [Figure S12B]. These tuning curves could be clustered into 7 tuning types [Figures 4C-D], and this analysis revealed that auditory responses are still divided into three main categories: pulse-tuned (tuning type 1), sine-tuned (tuning type 5-7), or non-selective (tuning type 2-4) [compare with Figure 3]. Sine-tuned ROIs were either low pass (tuning type 5-6: strongest responses to 100 Hz or 150 Hz sines, matching con-specific sine song frequencies) or had a broader distribution of preferred carrier frequencies (tuning type 7) [Figure 4Ei]. Pulse-tuned ROIs responded best to pulse frequencies, pauses and durations present in natural courtship song [Figure 4Eii]. Sine-tuned ROIs preferred continuous stimuli, which explains their preference for pulses with short pauses (below 20 ms) and longer pulse durations. Most tuning types were sensitive to the broad range of intensities tested (0.5-5 mm/s) with proportional increases in response with stimulus magnitude, and responded best to longer pulse train or sine tone durations (4s or longer) [Figures S12D-E]. Although, during natural courtship, pulse or sine trains (stretches of each type of song) have a mode of ∼360 ms and rarely last longer than 4s ^9^, preference for unnaturally long bouts have been recently reported ^34^. Finally, the distribution of tuning types shifted along the pathway from the AMMC to the six additional neuropils examined here [Figure 4D] – sine-tuned responses (tuning types 5-7) dominated in the regions of AMMC, SAD, and WED we imaged, while pulse-tuned responses (tuning type 1) dominated in the GNG, AVLP, and PVLP [Figure 4D]. In contrast, the PLP and LH regions we imaged contained an over-representation of sine-tuned responses (tuning types 5 and 6), accompanied by an increase in broadly tuned responses (tuning types 3-4). Not only do these results validate our choice of a single pulse and sine stimulus for probing the representation of song throughout the brain, they reveal how representations of courtship song are transformed across these neuropils.

**Figure 4.**
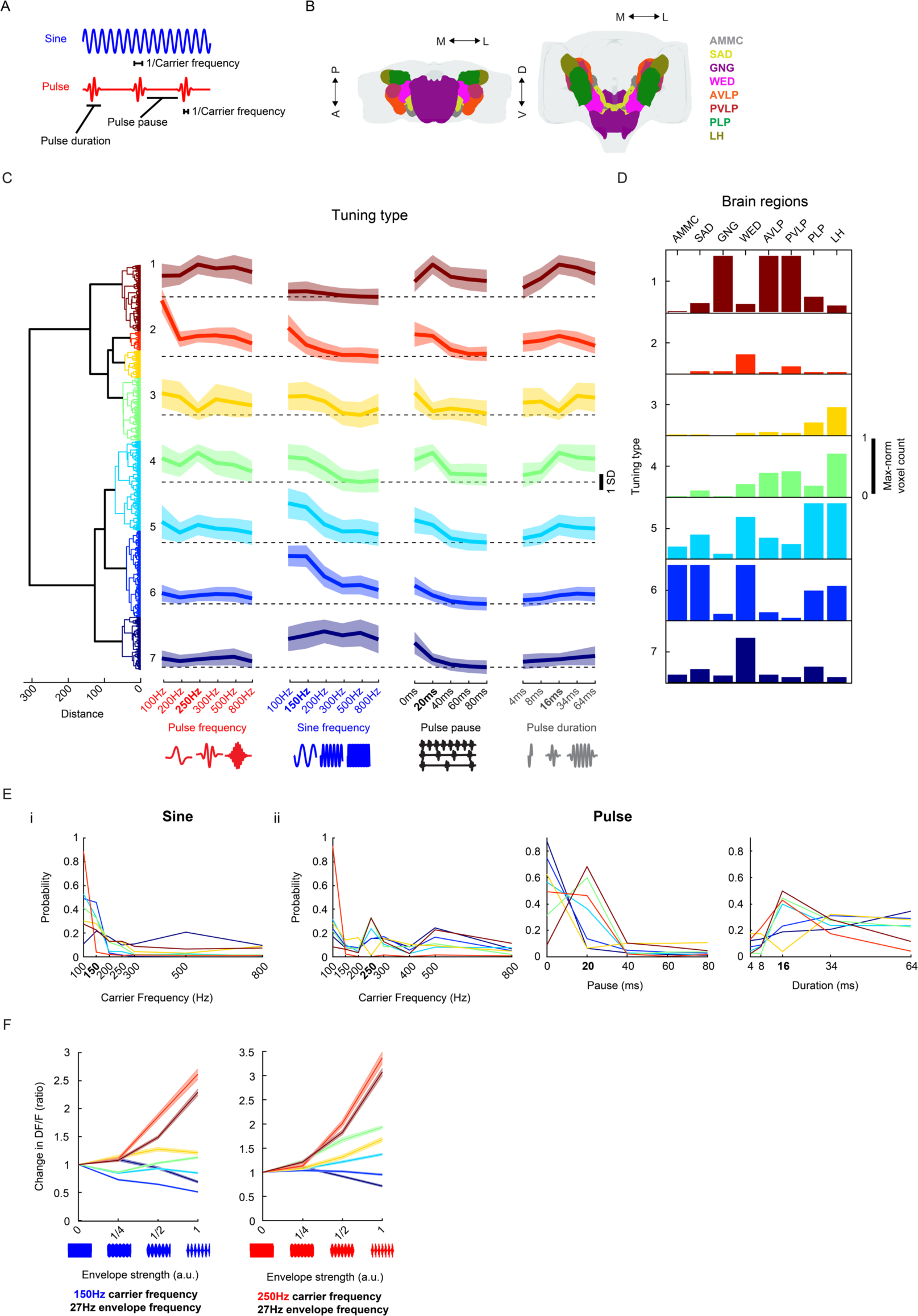
Widespread auditory activity is tuned to features of conspecific courtship songs. (A) Spectral and temporal features of the two main modes of *Drosophila* courtship song. (B) Mechanosensory neuropils imaged. (C) Hierarchical clustering of auditory tuning curves (see Figure S12A for how tuning curves were generated) into 7 distinct tuning types. Thick traces are the mean z-scored response magnitudes (80^th^ or 20^th^ percentile of activity - for excitatory or inhibitory responses - during stimuli plus 2 seconds after - activity is plotted as standard deviation (SD) of DF/F values) to the stimuli indicated on the X axis (across ROIs and within each tuning type) and shading is the standard deviation (n = 21 flies, 10,970 ROIs). Values of each feature for *Drosophila melanogaster* courtship song are indicated in bold on the X axis. (D) Distribution of tuning types across selected brain neuropils. Histogram of voxel count per type (number of voxels with activity that falls into each tuning type) is max-normalized for each neuropil (normalized per column). (E) Probability distributions of best frequency for sine (i) and pulse (ii), and best temporal feature (pulse duration and pause) for pulse (ii), separated by tuning type. Colorcode is the same as in (C). (F) Effect of amplitude envelope on responses to sine tones. Envelopes (∼27Hz envelope) of different amplitudes were added to sine tones (as depicted on the X axis, see Methods for more details) of 150 or 250 Hz. ROI response magnitudes to different envelope strengths were normalized to responses to envelope strength 0 (i.e., a sine tone of 150 or 250 Hz, with no envelope modulation) and sorted by tuning type from (C). Thick traces are the mean normalized response magnitudes per tuning type, and shading is the s.e.m. See also Figure S12.

**Figure 5.**
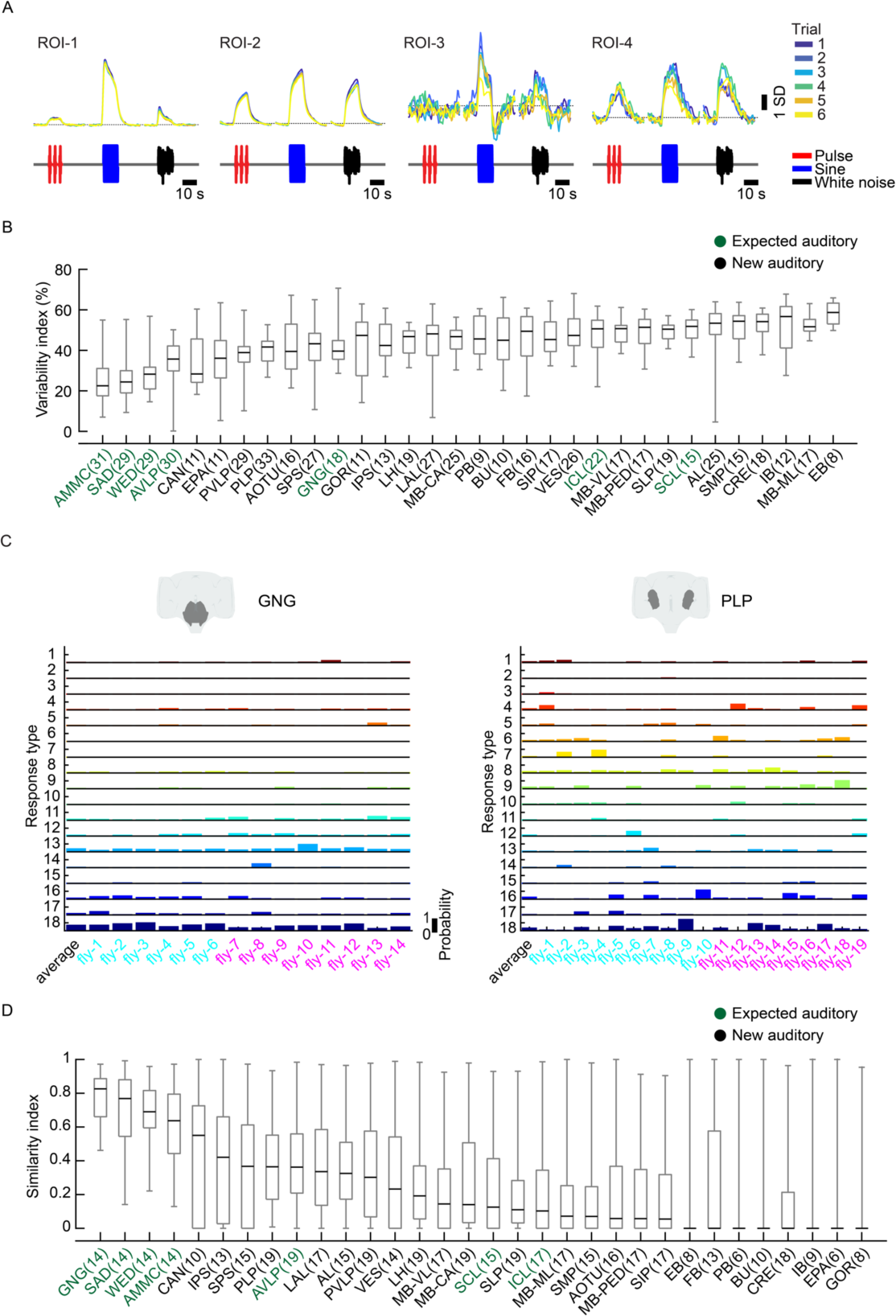
Auditory activity is more similar across trials and individuals in early mechanosensory areas. (A) Example auditory responses from five different ROIs to pulse, sine and white noise stimuli. Individual trials are plotted in different colors. All responses are z-scored, and therefore plotted as the standard deviation (SD) of the responses over time, black dash line corresponds to mean baseline across stimuli for each ROI. (B) Across-trial variability index of individual ROIs was computed as the residual of the variance explained by the mean across trials (see Methods). The grey box shows the 25^th^ and 75^th^ percentile; inner black line is the median variability across flies (number of flies per neuropil are shown in parentheses), whiskers correspond to minimum and maximum values. (C) Stereotypy of auditory response types (see Figures 3A-B) across individuals for the gnathal ganglion (GNG) and posterior lateral protocerebrum (PLP). Male flies indicated in cyan and female flies in magenta on X axis. Auditory activity in the GNG is similar across flies, while auditory activity in the PLP is more variable, in terms of response type diversity. Average response across all flies is shown in the left-most column. (D) Across-individual similarity index was computed by measuring the cosine between response type distributions (see Figure 3B) per neuropil across individuals (see Methods). Conventions same as in (B). Responses in the GNG, SAD, WED, and AMMC are the most stereotyped across individuals. See also Figures S13 and S14.

**Figure 6.**
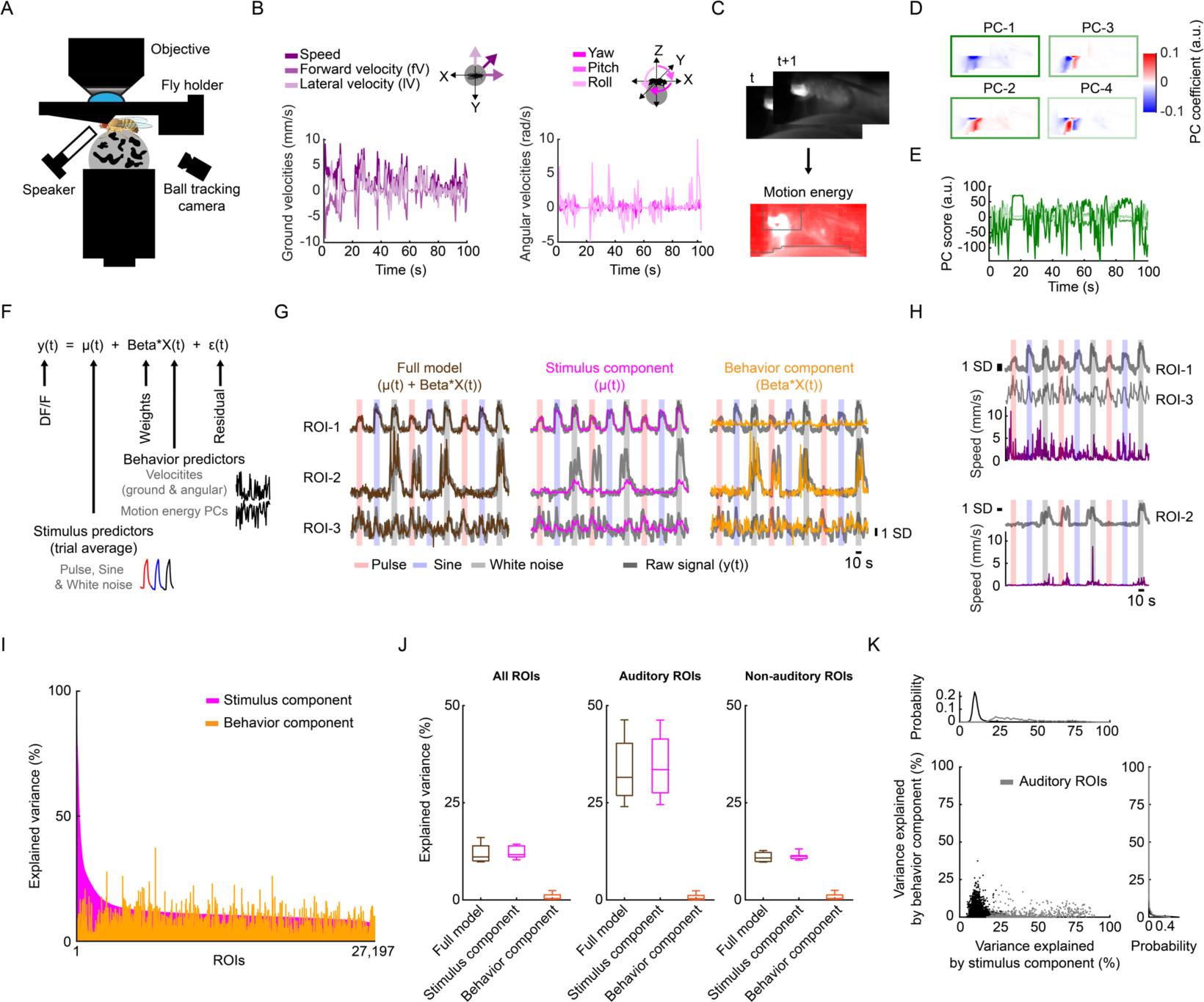
Spontaneous movements do not account for trial-to-trial variability in auditory responses. (A) Schematic of functional imaging of head-fixed fly walking on a ball. (B) Instantaneous ground and angular velocities measured from ball tracking of a representative tethered fly walking. (C) Motion energy (absolute value of the difference of consecutive frames) from a video recording of tethered fly. Grey contours correspond to pixels excluded from analysis (see Methods). (D) Coefficients of the top four PCs of the motion energy movie (image scale represents low-motion (blue) to high-motion energy (red)) of a representative tethered fly (same as (B)). (E) Scores of the top four PCs of the motion energy movie (Color of each PC is the same as in box color in D) of a representative tethered fly (same as (B)). (F) Schematic of linear model used to predict neural activity related to either stimuli or behavior. ROI’s activity (y(t)) is modelled as the sum of stimulus component (trial average, μ(t)), behavior component (Beta*X(t)), plus noise (ε(t)). (G) ROI activity along with predictions by stimulus component (μ(t)), behavior component (Beta*X(t)), or full model for three example ROIs most strongly modulated by auditory stimuli (ROI-1), by behavior (ROI-2), or by the combination of both (ROI-3). ROIs’ activity are z-scored, and DF/F units are in standard deviation (SD) (also for panel H). (H) Activity of ROIs from panel (G) along with speed. ROIs from the same fly are plotted together. (I) Explained variance by stimulus component or behavior component for individual ROIs, sorted by stimuli only model performance (n = 5, 27,197 ROIs). (J) Mean explained variance by stimulus component (μ(t)), behavior component (Beta*X(t)), or both (μ(t) + Beta*X(t)) across flies, for all ROIs (left), auditory ROIs only (middle), and non-auditory ROIs (right). The box shows the 25^th^ and 75^th^ percentile; inner line is the median explained variance across flies, whiskers correspond to minimum and maximum values. (K) Explained variance by stimulus component versus behavior component for individual ROIs. Grey dots correspond to auditory ROIs (using the same criteria as in Figure S1D). See also Figure S15.

*Drosophila melanogaster* pulse song is a sine tone with amplitude modulation at ∼27Hz, but responses to sine tones and pulses at various carrier frequencies correlated weakly across tuning types [Figure 4C]: responses tuned for pulse stimuli (e.g., tuning type 1) prefer pulses at 250Hz, but not 250Hz sines. This suggests that the slow frequency of the amplitude envelope has nonlinear interactions with carrier frequencies. In order to evaluate the categorical boundary between sine and pulse responses (in other words, the interaction between stimulus envelope and carrier frequency), we used a novel set of stimuli. We added a ∼27 Hz envelope to pure tones of 150 or 250 Hz carrier frequency at different strengths (see Methods). As expected, responses tuned to pulses increased their response with the strength of the envelope (in other words as a sine stimulus became more pulse-like), and the magnitude of this enhancement was carrier frequency dependent [Figure 4F]. Similarly, responses tuned to sine tones were strongly attenuated by the presence of an envelope, with different sensitivities by tuning type. Thus, we find a strong categorical boundary between pulse- and sine-tuned responses in the *Drosophila* brain, and little evidence for selectivity for intermediate envelopes and frequencies.

### Auditory activity ranges from stereotyped to variable across both trials and individuals

It is not known how reliable sensory responses are across trials, individuals, and sexes for most sensory modalities and brain areas. Our method of precise registration of volumetrically imaged auditory activity enabled us to systematically evaluate response variability across different neuropils and tracts. We first measured the residual of the variance explained by the mean across trials for all auditory ROIs (see Methods). Our criteria for selecting auditory ROIs [Figures S1D-F] excluded ROIs that did not show very consistent responses across trials to at least one auditory stimulus. Nonetheless, we still observed a range of across-trial variabilities for auditory ROIs [Figure 5A], with a pattern across neuropils: early mechanosensory neuropils (AMMC, SAD, WED, and AVLP) contained the lowest across-trial response variance, while all other neuropils exhibited higher trial-to-trial variability [Figure 5B]. This variation was, to some degree, correlated with the magnitude of auditory responses across neuropils [Figure S13A].

We showed above that particular response types [Figure 3A] are found in consistent spatial locations across flies [Figure 9B], suggesting a high degree of across-individual stereotypy of auditory responses. To quantify this, we used a similarity index (see Methods) to compare activity profiles across individuals [Figures 4C-D and S14]. Some neuropils, like the gnathal ganglion (GNG), contained ROIs with temporal responses highly similar across individuals (e.g., all individuals have a high frequency of response type 18 in the GNG), whereas other neuropils, like the posterior lateral protocerebrum (PLP), have more variable activity across individuals. Comparing the distribution of the similarity index across all neuropils revealed that early mechanosensory neuropils also have the highest across-individual similarity in auditory responses [Figures 4D and S13B]. Similarity decreases in downstream neuropils, and further decreases in neuropils outside the canonical mechanosensory system (e. g., visual, olfactory, and pre-motor neuropils). These differences in similarity across neuropils are robust to changes in how auditory responses are clustered [Figure S13C] and do not arise from variation in GCaMP fluorescence values across animals [Figure S13D].

### Spontaneous movements do not account for auditory response variability

Across-trial variability in auditory responses could arise from the time-varying behavioral state of the animal over the course of an experiment, as has been shown recently for cortical activity in mice ^35, 36^. Above, we did not track animal behavior, and so therefore could not correlate response variation with behavioral variation. To address this issue, we collected a new dataset in which we simultaneously recorded fly motion on a spherical treadmill while imaging pan-neuronally from sub-regions of the AMMC, SAD, AVLP, PVLP and LH [Figures 6A-E] – these regions were chosen because they contained a large number of auditory ROIs, with a range of across-trial variation from stereotyped (AMMC and SAD), to more variable (AVLP, PVLP, and LH) [Figure 4]. In these flies, we observed strong auditory activity, with across-trial variability similar to our previous dataset [Figure S15A]. Flies also showed a range of walking speeds on the ball [Figure S15B] that match natural speeds during fly courtship ^9^. Fly movements in these experiments appeared to be spontaneous because they were not reliably predicted by the auditory stimuli [Figure S15C]. In addition, despite observing auditory activity in pre-motor neuropils [Figure 3B], imaging from the fly ventral nerve cord in non-walking flies did not uncover much activity (only 247 out of 39,580 ROIs recorded from the ventral nerve cord were significantly correlated with any of the stimuli [Figures S15D-G]). This suggests that auditory ROIs in our study reflect sensory responses to stimuli, not motor responses.

Because flies moved spontaneously during presentation of auditory stimuli, this allowed us to determine what fraction of the across-trial variance in responses could be explained by the stimulus, by fly behavior, or by some combination of the two. We built a linear encoding model to predict neural responses [Figure 6F], similar to ^35^, from a combination of the mean response to the stimulus (µ(t)) and behavior predictors (X(t)), consisting of all tracked motor variables [Figures 6B-E] (see Methods). For this analysis we used all ROIs, not just those that passed our criteria for being ‘auditory’ [Figure S1D]. Of >27,000 ROIs segmented from the central brain, each had a different amount of explained variance from either the stimulus or behavior; some ROI responses tracked the stimulus [ROI-1, Figure 6G], while others tracked fly behavior [ROI-2, Figures 6G-H], and some ROI responses tracked both the stimulus and behavior [ROI-3, Figures 6G-H]. However, the stimulus component (µ(t)) explained the majority of variance, while the behavior component (Beta*X(t)) explained a much smaller fraction [Figure 6I] - the full model (µ(t) + Beta*X(t)) was as good at explaining the variance across ROIs as the stimulus component alone [Figure 6J]. When plotting the explained variance of the stimulus component against the behavior component, we found no significant trend for explained variance from the behavior predictors - that is, ROIs with little correlation to the stimulus had as much explained variance from the motor variables as ROIs with stronger correlation to the stimulus [Figure 6K]. These results suggest that the variability we observe across trials in auditory ROIs, at least for the brain regions imaged here, does not arise from fluctuations in spontaneous movements.

## DISCUSSION

Sensory systems are typically studied starting from the periphery and continuing to downstream partners guided by anatomy. This has limited our understanding of sensory processing to early stages of a given sensory pathway. Here, we used a brain-wide imaging method to unbiasedly screen for auditory responses beyond the periphery, and, via precise registration of recorded activity, to compare auditory representations across brain regions, individuals and sexes [Figure 1]. We found that auditory activity is widespread, extending well beyond the canonical mechanosensory pathway, and present in brain regions and tracts known to process other sensory modalities (i.e. olfaction and vision), or to drive motor behaviors [Figure 2]. The representation of auditory stimuli diversified, in terms of both temporal responses to stimuli and tuning for stimulus features, from the AMMC to later stages of the putative pathway, becoming more selective for particular aspects of courtship song (i.e. sine or pulse song, and their characteristic spectrotemporal patterns) [Figures 3-4]. Auditory representations were more stereotypic across trials and individuals in early stages of mechanosensory processing, and more variable at later stages [Figure 5]. By recording neural activity in behaving flies, we found that fly movements accounted for only a small fraction of the variance in neural activity, suggesting that across-trial auditory response variability stems from other sources [Figure 6]. These results have important implications for how the brain processes auditory information to extract salient features and to ultimately guide behavior.

Our understanding of the *Drosophila* auditory circuit thus far has been built up from targeted studies of neural cell types that innervate particular brain regions close to the auditory periphery ^14–18^. Altogether, these studies have delineated a pathway that starts in the Johnston’s Organ (JO), and extends from the antennal mechanosensory and motor center (AMMC), to the wedge (WED), ventrolateral protocerebrum (VLP), and lateral protocerebral complex (LPC). By imaging pan-neuronally, we found widespread auditory responses, spanning brain regions beyond the canonical pathway, suggesting auditory processing is more distributed. However, for neuropil signals it is challenging to determine the number of neurons that contribute to the ROI responses we describe. Although the diverse set of temporal and tuning types per neuropil [Figure 3] suggests we have sampled many neurons per neuropil, restricting GCaMP to spatially-restricted genetic enhancer lines [Figures S6C-D], will assist with linking broad functional maps with the cell-types constituting them.

Our findings of widespread auditory activity are likely not unique to audition. So far, in adult *Drosophila*, only taste processing has been surveyed brain-wide ^37^. While that study did not map activity onto neuropils and tracts, nor made comparisons across individuals, it suggested that taste processing was distributed throughout the brain. Similarly in vertebrates, widespread responses to visual, and nociceptive have been observed throughout the brain ^38, 39^. Our findings are consistent with anatomical studies that find connections from the AMMC and WED to several other brain regions ^18, 40^, and with our own analysis of the ‘hemibrain’ connectome [Figure 2]. While we do not yet know what role this widespread auditory activity plays in behavior, we show that ROIs that respond to auditory stimuli do so with mostly excitatory (positive-going) responses and that activity throughout the brain is predominantly tuned to features of the courtship song. During courtship, flies evaluate multiple sensory cues (olfactory, auditory, gustatory and visual) to inform mating decisions and modulate their mating drive. Although integration of multiple sensory modalities have been described in higher order brain regions ^41^, our results suggest that song representations are integrated with olfactory and visual information at earlier stages [Figures 2-3]. In addition, song information may modulate the processing of non-courtship stimuli. Song representations in the mushroom body may be useful for learning associations between song and olfactory, gustatory, or visual cues ^42^, while diverse auditory activity throughout all regions of the lateral horn may indicate an interaction between song processing and innate olfactory behaviors ^43^. Finally, we found auditory activity in brain regions involved in locomotion and navigation (central and lateral complex, superior and ventromedial neuropils). Activity in these regions is quite diverse [Figure 3], suggesting that pre-motor circuits receive information about courtship song patterns, and could therefore underlie stimulus-specific locomotor responses ^4^.

*Drosophila melanogaster* songs are composed of pulses and sines, which differ in their spectral and temporal properties - it is unclear how and where selectivity for the different song modes arises in the brain. Since neurons in the lateral protocerebral complex (LPC) are tuned for pulse song across all timescales that define that mode of song ^4^, neurons upstream must carry the relevant information to generate such tuning. Here we find many ROIs that are selective for either sine or pulse stimuli throughout the entire central brain [Figure 3]. More systematic examination of tuning in the neuropils that carry the most auditory activity (antennal mechanosensory and motor center (AMMC), saddle (SAD), wedge (WED), anterior and posterior ventrolateral protocerebrum (AVLP and PVLP), posterior lateral protocerebrum (PLP), and lateral horn (LH)) revealed that most ROIs are tuned to either pulse or sine features, with few ROIs possessing intermediate tuning [Figure 4]. This suggests that sine and pulse information splits early in the pathway. We also found that sine-selective responses dominate throughout the brain - while some of this selectivity may simply reflect preference for continuous versus pulsatile stimuli, our investigation of feature tuning revealed that many of these ROIs preferred frequencies present specifically in courtship songs [Figure 4]. Previous studies indicated that pulse song is more important for mating decisions, with sine song purported to play a role in only priming females ^5, 44^. However our results, in combination with the fact that males spend a greater proportion of time in courtship singing sine versus pulse song ^9^, suggest a reevaluation of the importance and role of sine song in mating decisions. This study therefore lays the foundation for exploring how song selectivity arises in the brain.

Our results also revealed that early mechanosensory brain areas contain ROIs with auditory activity less variable across trials and animals [Figure 5]. Our results for across-animal variability have parallels to the *Drosophila* olfactory pathway: 3^rd^ order mushroom body neurons are not stereotyped, while presynaptic neurons in the antennal lobe, the projection neurons, are ^45^. Similarly, a lack of stereotypy beyond early mechanosensory brain areas may reflect stochasticity in synaptic wiring. The amount of variation we observed in some brain areas is quite large [Figure S14], and follow up experiments with sparser driver lines will be needed to validate whether what we report here applies to variation across individual identifiable neurons. We also observed a wide range of across-trial variability throughout neuropils with auditory activity [Figure 5B]. Imaging from a subset of brain regions in behaving flies revealed that trial-to-trial variance in auditory responses is not explained by spontaneous movements [Figure 6], suggesting variance is driven by internal dynamics. This result differs from recent findings in the mouse brain which showed that a large fraction of activity in sensory cortices corresponds to non-task-related or spontaneous movements ^35, 36^ – this may indicate an important difference between invertebrate and vertebrate brains, and the degree to which ongoing movements shape activity across different brains. However, we should point out that motor activity is known to affect sensory activity in flies ^46^, but this modulation is tied to movements that are informative for either optomotor responses or steering ^47, 48^. In our experiments, flies were not behaviorally engaged with the auditory stimuli – while playback of the same auditory stimuli can reliably change walking speed in freely behaving flies ^4^, we did not observe reliable behavioral responses here. Adjusting our paradigm to drive such responses ^49^, might uncover behavioral modulation of auditory activity. Alternatively, behavioral modulation of auditory responses may occur primarily in motor areas, such as the central complex or areas containing projections of descending neurons ^50^. Further dissection of the sources of this variability would require capturing more brain activity simultaneously in behaving animals, while not significantly compromising spatial resolution.

Just as the discovery of movement-related activity throughout the brain ^21, 35, 36, 51^ motivates fine-scale tracking of animal behavior during recordings, regardless of task or paradigm used, our discovery of widespread auditory activity throughout the fly central brain motivates the use of methods to systematically record activity across the brain when studying coding of any sensory modality. Here we provide such tools for characterizing sensory activity all registered in the same atlas coordinates for comparisons across trials, individuals, and sexes. By producing maps for additional modalities and stimulus combinations, and by combining these maps with information on connectivity between and within brain regions ^31, 52^, the logic of how the brain represents the myriad stimuli and their combinations present in the world will emerge.

## SUPPLEMENTAL INFORMATION

Supplemental information includes 15 figures and 5 movies.

## ACKNOWLEDGEMENTS

We thank Jan Clemens for assistance on auditory stimuli delivery and linear modeling of calcium responses, and helpful discussions on auditory coding, Andrea Giovannucci for help with ROI segmentation of volumetric calcium signals and guidance with CaImAn toolbox usage, Gregory Jefferis and Torsten Rohlfing for help using the image registration toolbox CMTK and the neuroanatomy toolboxes NAT and NBLAST, Ben Cowley for help with linear modeling of calcium responses in behaving flies, Adam Calhoun for help with analysis of the ‘hemibrain’ connectome dataset, and Talmo Pereira for assistance with the Coherent Point Drift algorithm. We also thank Greg Jefferis, Sama Ahmed, Christa Baker, and Jan Clemens for comments on the manuscript. DAP was supported in part by an NSF Physics Frontier Center grant, and MM was supported by an NIH NINDS New Innovator award, NIH BRAIN Initiative R01s NS104899 and NS110060, and a Howard Hughes Medical Institute Faculty Scholar award.

## AUTHOR CONTRIBUTIONS

D.A.P. and M.M. designed the study. D.A.P. collected and analyzed data. ST designed and built the two-photon imaging microscope. E.P. generalized ROI segmentation from 2D to 3D datasets. D.A.P. and M.M. wrote the manuscript.

## DECLARATION OF INTERESTS

The authors declare no competing interests.

## DATA AND CODE AVAILABILITY

All calcium imaging data from this study (segmented ROIs) is publicly available at xx[DP1]. All code for the neural imaging data processing pipeline is available at https://github.com/dpacheco0921/CaImProPi[DP2].

## KEY RESOURCES TABLE

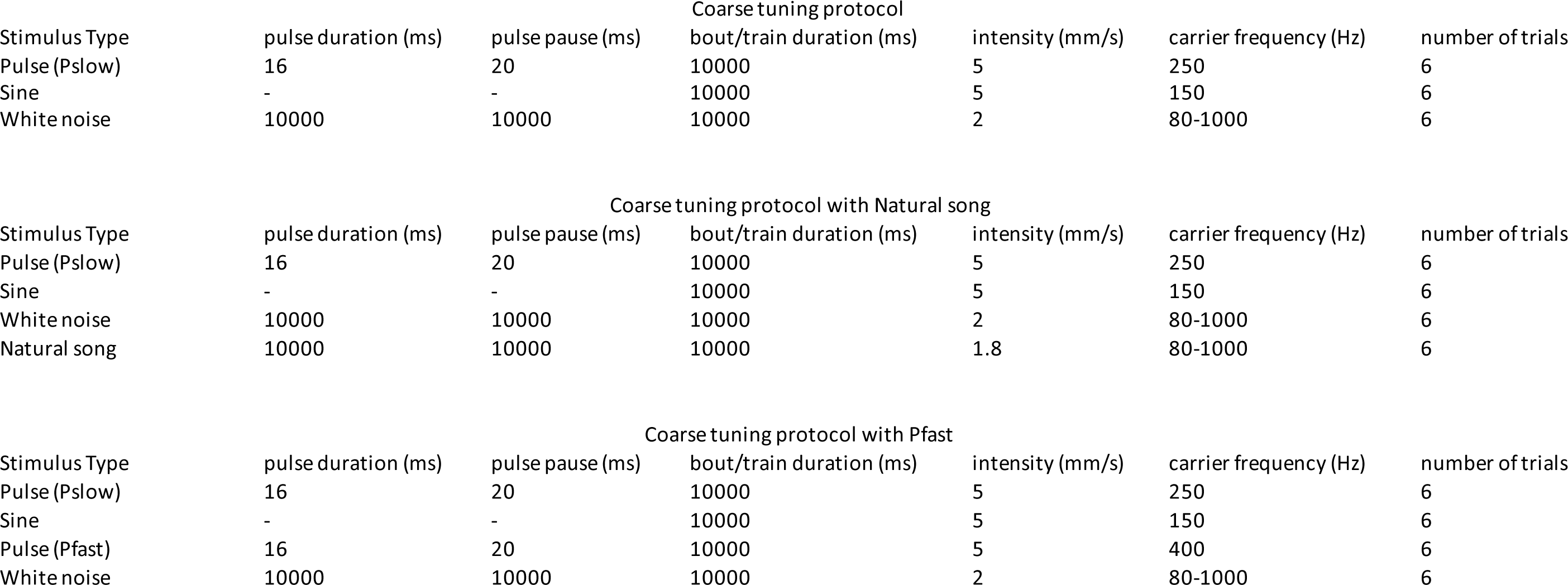

| Stimulus Type | Coarse tuning protocol |  |  |  |  |  |
| --- | --- | --- | --- | --- | --- | --- |
|  | pulse duration (ms) | pulse pause (ms) | bout/train duration (ms) | intensity (mm/s) | carrier frequency (Hz) | number of trials |
| Pulse (Pslow) | 16 | 20 | 10000 | 5 | 250 | 6 |
| Sine | - | - | 10000 | 5 | 150 | 6 |
| White noise | 10000 | 10000 | 10000 | 2 | 80-1000 | 6 |

| Stimulus Type | Coarse tuning protocol with Natural song |  |  |  |  |  |
| --- | --- | --- | --- | --- | --- | --- |
|  | pulse duration (ms) | pulse pause (ms) | bout/train duration (ms) | intensity (mm/s) | carrier frequency (Hz) | number of trials |
| Pulse (Pslow) | 16 | 20 | 10000 | 5 | 250 | 6 |
| Sine | - | - | 10000 | 5 | 150 | 6 |
| White noise | 10000 | 10000 | 10000 | 2 | 80-1000 | 6 |
| Natural song | 10000 | 10000 | 10000 | 1.8 | 80-1000 | 6 |

| Stimulus Type | Coarse tuning protocol with Pfast |  |  |  |  |  |
| --- | --- | --- | --- | --- | --- | --- |
|  | pulse duration (ms) | pulse pause (ms) | bout/train duration (ms) | intensity (mm/s) | carrier frequency (Hz) | number of trials |
| Pulse (Pslow) | 16 | 20 | 10000 | 5 | 250 | 6 |
| Sine | - | - | 10000 | 5 | 150 | 6 |
| Pulse (Pfast) | 16 | 20 | 10000 | 5 | 400 | 6 |
| White noise | 10000 | 10000 | 10000 | 2 | 80-1000 | 6 |

| Stimulus Type | pulse duration (ms) | pulse pause (ms) | bout/train duration (ms) | Feature tuning protocol |  | number of trials | envelope strength | envelope frequency (Hz) |
| --- | --- | --- | --- | --- | --- | --- | --- | --- |
|  |  |  |  | intensity (mm/s) | carrier frequency (Hz) |  |  |  |
| Pulse | 16 | 20 | 4000 | 5 | 250 | 6 | - | - |
| Sine | - | - | 4000 | 5 | 150 | 6 | - | - |
| White noise | 10000 | 10000 | 4000 | 2 | 80-1000 | 6 | - | - |
| Pulse carrier | 16 | 20 | 4000 | 5 | 100 | 3 | - | - |
| Pulse carrier | 16 | 20 | 4000 | 5 | 150 | 3 | - | - |
| Pulse carrier | 16 | 20 | 4000 | 5 | 200 | 3 | - | - |
| Pulse carrier | 16 | 20 | 4000 | 5 | 300 | 3 | - | - |
| Pulse carrier | 16 | 20 | 4000 | 5 | 500 | 3 | - | - |
| Pulse carrier | 16 | 20 | 4000 | 5 | 800 | 3 | - | - |
| Sine carrier | - | - | 4000 | 5 | 100 | 3 | - | - |
| Sine carrier | - | - | 4000 | 5 | 200 | 3 | - | - |
| Sine carrier | - | - | 4000 | 5 | 250 | 3 | - | - |
| Sine carrier | - | - | 4000 | 5 | 300 | 3 | - | - |
| Sine carrier | - | - | 4000 | 5 | 500 | 3 | - | - |
| Sine carrier | - | - | 4000 | 5 | 800 | 3 | - | - |
| Pulse pause | 16 | 0 | 4000 | 5 | 250 | 3 | - | - |
| Pulse pause | 16 | 40 | 4000 | 5 | 250 | 3 | - | - |
| Pulse pause | 16 | 60 | 4000 | 5 | 250 | 3 | - | - |
| Pulse pause | 16 | 80 | 4000 | 5 | 250 | 3 | - | - |
| Pulse duration | 4 | 20 | 4000 | 5 | 250 | 3 | - | - |
| Pulse duration | 8 | 20 | 4000 | 5 | 250 | 3 | - | - |
| Pulse duration | 32 | 20 | 4000 | 5 | 250 | 3 | - | - |
| Pulse duration | 64 | 20 | 4000 | 5 | 250 | 3 | - | - |
| Pulse intensity | 16 | 20 | 4000 | 0.5 | 250 | 3 | - | - |
| Pulse intensity | 16 | 20 | 4000 | 1 | 250 | 3 | - | - |
| Pulse intensity | 16 | 20 | 4000 | 2 | 250 | 3 | - | - |
| Sine intensity | - | - | 4000 | 0.5 | 150 | 3 | - | - |
| Sine intensity | - | - | 4000 | 1 | 150 | 3 | - | - |
| Sine intensity | - | - | 4000 | 2 | 150 | 3 | - | - |
| Pulse train duration | 16 | 20 | 2000 | 5 | 250 | 3 | - | - |
| Pulse train duration | 16 | 20 | 8000 | 5 | 250 | 3 | - | - |
| Sine bout duration | - | - | 2000 | 5 | 150 | 3 | - | - |
| Sine bout duration | - | - | 8000 | 5 | 150 | 3 | - | - |
| Sine plus envelope | - | - | 4000 | 5 | 150 | 3 | 0 | 30.3 |
| Sine plus envelope | - | - | 4000 | 5 | 150 | 3 | 0.25 | 30.3 |
| Sine plus envelope | - | - | 4000 | 5 | 150 | 3 | 0.5 | 30.3 |
| Sine plus envelope | - | - | 4000 | 5 | 150 | 3 | 1 | 30.3 |
| Sine plus envelope | - | - | 4000 | 5 | 150 | 3 | 0 | 27.8 |
| Sine plus envelope | - | - | 4000 | 5 | 150 | 3 | 0.25 | 27.8 |
| Sine plus envelope | - | - | 4000 | 5 | 150 | 3 | 0.5 | 27.8 |
| Sine plus envelope | - | - | 4000 | 5 | 150 | 3 | 1 | 27.8 |

## CONTACT FOR REAGENT AND RESOURCE SHARING

Further information and requests for resources and reagents should be directed to the Lead Contact Mala Murthy

## EXPERIMENTAL MODEL AND SUBJECT DETAILS

### Fly Stocks

Flies used to generate the IVIA atlas and for functional imaging experiments were of the following genotype: w/+; GMR57C10-LexA/+; 13xLexAop-GCaMP6s, 8xLexAop-mCD8tdTomato/+ (nsyb-LexA-G6s-tdtom), all ‘+’ chromosomes came from the NM91 wild type strain ^1^. *iav*^1^ mutants had the following genotype: *iav*^1^/Y; GMR57C10-LexA/+; 13xLexAop-GCaMP6s, 8xLexAop-mCD8tdTomato/+. For imaging of olfactory projection neurons and Kenyon cells, flies had the following genotype: w; GMR57C10-LexA, 8xLexAop-myr-tdTomato/GH146-Gal4; 20xUAS-GCaMP6s/+ and w; GMR57C10-LexA, 8xLexAop-myr-tdTomato/+; 20xUAS-GCaMP6s/OK107, ey respectively. For stalk segmentation of pC1, pC2l and pC2m Dsx+ neurons w; GMR57C10-LexA/20xUAS-GCaMP6s, 20xUAS-myr-tdTomato; 8xLexAop-mCD8tdTomato/dsx^Gal4^, for stalk segmentation of P1 neurons and pMP7 neurons: w; NP2631-Gal4, 20XUAS>STOP>CsChrimson.mVenus/GMR57C10-LexA; 8xLexAop-mCD8tdTomato/ fruFLP. We acquired 13xLexAop-GCaMP6s (44590), GH146-Gal4(30026), and OK107 (854) from the Bloomington stock center, iav^1^ from the Kyoto stock center, GMR57C10-LexA from Gerry Rubin, 20XUAS>STOP>CsChrimson.mVenus from Vivek Jayaraman, 8xLexAop-mCD8tdTomato from Yuh Nung Jan, NP2631-Gal4 from the Kyoto stock center, fru^FLP^ from Berry Dickson, dsx^Gal4^ from Stephen Goodwin, NM91 from the Andolfatto group at Columbia University, and 20xUAS-myr-tdTomato from Glenn Turner.

## METHODS

### Sound delivery and acoustic stimuli

Sound was delivered as in^2^. During experiments, the sound tube was positioned in front of the fly at a distance of 2 mm and an angle of ∼17⍛ from the midline (elevation of ∼39⍛). In this configuration, we observed qualitatively similar responses from the AMMC in both hemispheres.

Acoustic stimuli were organized in two main protocols: a) coarse tuning protocol, and b) feature tuning protocol (see supplemental Table S2 for a list of all stimulus sets used). For both protocols the intensity of auditory stimuli presented was within the dynamic range of JO neurons ^3^ (0.1-6 mm/s) and within the distribution of song intensities produced during natural courtship ^4^ (pulse and sine song intensity 5mm/s, white noise intensity 2 mm/s). Stimuli were preceded and followed by 10 or 4 seconds of silence, for coarse tuning and feature tuning protocol respectively, to reduce crosstalk between responses to subsequent stimulus. Stimuli was presented in blocks (e.g. pulse, sine, white noise repeated 6 times) or randomized for coarse tuning and feature tuning protocol respectively. We chose long stimuli duration (10 or 4 seconds) to maximize signal-to-noise ratio of auditory responses.

### Head-fixed walking setup and behavioral tracking

We shaped 9 mm diameter balls from polyurethane foam (FR-7120, General Plastics) similar to ^5^. The ball rested in an aluminium ball holder with a concave hemisphere 9.6 mm in diameter [Figure 6A]. The ball holder had a 1.27 mm channel drilled through the bottom of the hemisphere and connected to air flowing at ∼75 ml/min. The ball holder was mounted on a manipulator (MS3, Thorlabs) to adjust the position of the ball under each fly.

We imaged the fly and ball from the side (90⍛ to the anterior-posterior axis of the fly) illuminated by a pair of IR 850 nm LEDs (M850F2, Thorlabs) - coupled to optic fibers and collimator lenses (F810SMA-780, Thorlabs) - imaging was done with a monochromatic camera (Flea3 FL3-U3-13Y3M-C ½, FLIR) externally triggered at 100 Hz, with a zoom lens (C-Mount 15.5-20.4mm Varifocal, Computar). The lens had a 770-790 nm narrow band filter (BN785, MidOpt) attached to block two-photon excitation laser (920 nm). This camera was used to position the fly and to track both the ball and fly movement.

To track the ball rotation we used a windows implementation of FicTrac software ^6^ (see https://github.com/murthylab/fic-trac-win). FicTrac calculates the angular position of the ball for each frame and reconstructs the X and Y trajectory on a fictive 2D surface. The ball was marked with non-repetitive shapes using black permanent marker (PK10, SARSTEDT) to allow tracking of the ball’s rotation.

We used the X, and Y trajectories to calculate instantaneous ground velocities (forward and lateral velocity, and the speed in the fictive heading direction (forward plus lateral velocity)), and the angular position to calculate the instantaneous angular velocities (yaw, pitch, and roll) [Figure 6B].

To extract a high-dimensional representation of the fly’s body motion (i.e. legs, abdomen and wings) we used the toolbox facemap ^7^ (see https://github.com/MouseLand/facemap). Facemap applies singular value decomposition to the motion energy movie (the absolute difference between frame_t_ and frame_t+1_) [Figure 6C], extracting the top 500 components’ coefficients (spatial weights) and corresponding scores (temporal profiles) of the motion energy movie [Figures 6D-E]. For this analysis the field of view of each recorded video was reduced to the smallest box containing the whole fly body, masking out pixels that belonged to the ball, and head plus anterior thorax (pixels of these body parts were removed due to its contamination with imaging related signal). The top 500 components of the motion energy movie per fly were used as a summary of motor behaviors for later analysis (see Linear modelling of neural activity from stimulus and behavior).

### Fly preparation and functional Imaging

Virgin female or male flies (3-7 days old) were mounted and dissected as described previously ^8, 9^, with minor differences, for non-behaving animals. The angle of the thorax to the posterior side of the head was kept close to 90°, keeping the head as parallel as possible to the holder floor. Following dissection of the head cuticle, we removed the air sacks and tracheas using sharp forceps (Dumont #5SF) and any additional fat or soft tissue was removed with suction using a sharp glass pipette. In addition, to enable imaging of ventral central brain regions close to the neck, we pushed the thorax posteriorly and fixed it using a tissue adhesive (3M vetbond) delivered using a sharp glass pipette. Proboscis or digestion-related motion artifacts were minimized by pulling the proboscis and waxing it at an extended configuration and removing muscle M16. In addition, during dissection we kept the antennae dry and mobile for auditory stimulation. For VNC recordings, in addition to the previous steps, we removed thoracic tissue dorsal to the VNC (e.g. cuticle, indirect flight muscles, and digestive system), exposing the 1^st^ and 2^nd^ segments of the VNC. For head-fixed behaving animals, flies were fixed to a custom holder (similar to ^10^) by gluing the lateral and dorsal anterior edges of the thorax with light cured glue (Bondic), followed by waxing of the posterior side of the head (the head was pushed forward to end up with a similar configuration as in dissection for non-behaving animals). The rest of the dissection was the same as for non-behaving animals.

Imaging experiments were performed on a custom-built two-photon laser scanning microscope equipped with 5 mm galvanometer mirrors (Cambridge Technology), an electro-optic modulator (M350-80LA-02 KD*P, Conoptics) to control the laser intensity, a piezoelectric focusing device (P-725, Physik Instrumente), a Chameleon Ultra II Ti:Sapphire laser (Coherent) and a water immersion objective (Olympus XLPlan 25X, NA=1.05). The fluorescence signal collected by the objective was reflected by a dichroic mirror (FF685 Dio2, Semrock), filtered using a multiphoton short-pass emission filter (FF01-680/sp-25, Semrock), split by a dichroic mirror (FF555 Dio3, Semrock) into two channels, green (FF02-525/40-25, Semrock) and red (FF01-593/40-25, Semrock), and detected by GaAsP photo-multiplier tubes (H10770PA-40, Hamamatsu). We used a laser power below 20mW, power above this level induced global increases in fluorescence. The microscope was controlled in Matlab using ScanImage 5.1 (Vidrio). Dissection chambers were placed beneath the objective, and perfusion saline was continuously delivered directly to the meniscus. The temperature of the perfusion saline was kept at 24° C using a miniature perfusion cooler and heater unit (TC-RD, TC2-80-150-C, Biosciencetools). We chose GCaMP6s over faster sensors to maximize signal-to-noise ratio of auditory responses. For wild type flies, we only recorded from animals that first exhibited auditory-evoked responses in the AMMC of both hemispheres. Recordings typically lasted for 2-3 hours. Flies whose global baseline fluorescence saturated and became homogeneous were discarded.

### Functional Imaging Analysis Pipeline

All steps from image preprocessing (I) to registration of functional data to IVIA (IV) are schematized in Supplemental Figure 1A, in which we take one example dataset step-by-step through the entire functional imaging analysis pipeline.

#### I. Image preprocessing

For all experiments, we recorded data from each animal using either of the following protocols: a) 14 to 17 subvolumes at 1 Hz until covering the whole extent of the Z axis per fly (posterior to anterior) - 1.2×1.4×2 µm^3^ voxel size and ∼542 kHz pixel rate [Figures 1-3, 5, S5C-E, S5G-H, S6C-D, and S15D-G]. The full volume scanned per fly was ¼ of the central brain or ∼300x300x250 µm^3^; b) 4 to 6 subvolumes from selected brain regions (not necessarily contiguous) at 2Hz - with 1x1x2 µm^3^, 1x1x3 µm^3^ or 1.2×1.2×5 µm^3^ voxel size and ∼542 kHz pixel rate [Figure 4]; c) One subvolume that extends from the lateral horn (LH) in the posterior brain to the antennal mechanosensory and motor center (AMMC) in the anterior brain at 1Hz - 1.2×1.4×15 µm^3^ voxel size and ∼542 kHz pixel rate [Figure 6]. To generate the private whole brain atlas (per fly), we imaged 1-4 Z-stack volumes (in both GCaMP and tdTomato channels) with 0.75×0.75×1 µm^3^ voxel size to span the entire central brain (both hemispheres) in the Z dimension.

For each volumetric time series or segment (XYZ and Time) - see Figure S1A - we performed rigid motion correction on the X, Y, and Z axes on the tdTomato signal using the NoRMCorre algorithm ^11^, and this transformation was then applied to the GCaMP6s signal. Segments with anisotropic XY voxel size were spatially resampled to have isotropic XY voxel size (bilinear interpolation on X and Y axes), and temporally resampled to correct for different slice timing across planes of the same volume. Finally, segments were up-sampled (linear interpolation) to twice the original sampling rate (2 Hz and 4 Hz, for segments recorded at 1 Hz and 2 Hz respectively), and aligned relative to the start of the first stimulus (linear interpolation) prior to analysis.

Segments consecutively recorded along the Z axis (for protocol a)) were stitched along the Z axis using NoRMCorre (similarly, we used the average tdTomato signal as the reference image and apply the transformation to the GCAMP6s signal) obtaining a ‘volume’. For protocols b) and c) each segment is treated as an independent ‘volume’. Volumes were then mirrored to the right hemisphere. Average tdTomato images were saved as individual NRRD (http://teem.sourceforge.net/nrrd) files for registration to IVIA (using Matlab, nrrdWriter function).

For the private whole brain atlases (per fly), Z-stacks (∼384×384×280 µm^3^ each) were stitched on the X and Y axes using a pairwise stitching algorithm in FIJI ^12, 13^. These whole brain images were smoothed on the X and Y axes (Gaussian kernel size of [3, 3] voxels and gaussian kernel standard deviation of [2, 2]), mirrored on the X axis as for the volumetric time series. Images were split into tdTomato and GCaMP6s channels and saved separately as NRRD files for registration to IVIA (using MatLab, nrrdWriter function).

#### II. ROI segmentation

We used the constrained non-negative matrix factorization (CNMF) algorithm ^14^ for ROI segmentation. The algorithm was implemented and generalized to 3D-data in the MATLAB version of the CaImAn toolbox ^15^. We used the greedy initialization (Gaussian kernel size of [9, 9, 5] voxels and gaussian kernel standard deviation of [4, 4, 2], and 10^th^ percentile baseline subtraction) with 600 components per sub-stack of 11 planes and walk through the entirety of the Z axis with a sub-stack overlap of 3 planes. After initialization, 1) spatial components were updated using the ‘dilate’ search method (the ‘ellipse’ method imposes a prior on the shape which is not optimal for detecting ROIs in neuropil), and default settings were used for updating temporal components, 2) spatially overlapping components were iteratively merged if the Pearson correlation of temporal components between them was greater than 0.9 (this allows stitching of contiguous patches of neuropil with similar temporal profiles). The combination of these two steps segmented soma-like and neuropil-like ROIs accordingly. We wrote custom code to compile spatial and temporal components from all sub-stacks per fly (similar to run_CNMF_patches.m), and to calculate DF(t)/Fo(t) ROI. Background fluorescence Fo(t) and delta-fluorescence DF(t) (i.e. ROI activity) per ROI are directly modelled by CaImAn, and provided as output variables. To calculate the DF(t)/Fo(t), first, for each ROI, we detrended DF(t) (we perform a 20^th^ percentile filtering over time, calculated from overlapping windows of 60 seconds each). Second, for each ROI we added to Fo(t) its corresponding trend removed from DF(t). DF(t)/Fo(t) was calculated as the detrended DF(t) divided by Fo(t) (which is now background fluorescence plus trend from DF(t)) for each time point. See https://github.com/dpacheco0921/CaImProPi.

#### III. Identification of stimulus-modulated ROIs

We linearly modelled each ROI signal as a convolution of the stimuli history and a set of three filters (one per stimulus) [Figure S1D]. Filters were estimated using ridge regression ^16^, with a filter length of 10 and 6 seconds for the coarse tuning protocol (Figures 1-3 and 5) and the feature tuning protocol (Figure 4), respectively. For filter estimation, we partitioned each ROI raw signal (F(t)) into training (80%) and testing (20%) datasets. Estimated filters (q(Τ)) were then convolved with the stimulus history (f(t)) to generate the predicted signal (g(t)) for each ROI. Prediction goodness was measured as the Pearson correlation coefficient between raw and predicted signals ρ(F, g) using fifthteen fold cross-validation. Statistical significance of correlation coefficients (ρ(F, g)) was determined by bootstrapping. Each ROI raw signal was randomly shuffled in chunks of 10 seconds (sF(t)), and a distribution of 10,000 correlation coefficients (ρ(sF, g))) between each independent shuffle and the predicted signal was generated. p-values for ρ(F, g) significance were calculated as the fraction of ρ(sF, g) with values greater than the 30th percentile of ρ(F, g). To correct for multiple comparisons of all ROIs within a fly, p-values were corrected using the Benjamini-Hochberg (B-H) FDR correction algorithm (FDR = 0.01). This correction assumes independence or positive dependency between tests.

#### IV. Registration of functional data to IVIA

*In vivo* functional data from each fly was registered to the IVIA using the structural channel (mCD8-tdTomato signal) in a two-step fashion: i) ‘volumes’ to private whole brain registration and ii) private whole brain to IVIA atlas registration [Figure S1A].

i) ‘volume’ to private whole brain registration: for each fly, we registered (linear followed by nonlinear registration) the volume tdTomato image to its own whole brain tdTomato image (Z-stack volume, see Image preprocessing). Given that the images come from the same individuals, we expected minimal nonlinear deformations. Therefore, we limited the linear transformation to only include rotation, translation and anisotropic scaling (metric: normalized mutual information). For the nonlinear transformation (metric: normalized mutual information), we used a small grid spacing (20 µm, with 2 refinements), Jacobian constraint weight of 0.01, and smoothness constraint weight of 0.1.

ii) Private whole brain to IVIA atlas registration: for each fly, we registered (linear followed by nonlinear registration) the private whole brain tdTomato image to the IVIA atlas. In this case the linear transformation included rotation, translation, anisotropic scaling, and shearing (metric: normalized mutual information). For the nonlinear transformation (metric: normalized mutual information), we used a bigger grid spacing (170 µm, with 5 refinements), Jacobian constraint weight of 0.001, and smoothness constraint weight of 1. We excluded the voxels covering the optic lobes for the calculation of the linear transformation, as this provided a better initialization for the nonlinear registration calculated using all voxels. This modification improved the registration success, and the quality of registration for central brain structures (visually inspected).

These i) volume to whole brain and ii) whole brain to IVIA atlas transformations per fly were concatenated to transform spatial components of segmented ROIs from the native volume coordinates to *in vivo* atlas coordinates.

### Construction of average in vivo intersex atlas (IVIA)

Recently, an in vivo D. melanogaster female atlas was built ^17^, however it does not account for known anatomical differences between female and male brains ^18^. Therefore, we built an average *in vivo* intersex brain atlas using a pool of 5 male and 8 female brains (expressing membranal tdTomato pan-neuronally (mCD8-tdTomato)) [Figure S2A]. In the first iteration, we picked a seed brain, half cropped on the X axis, and stitched it to its mirror image to generate a symmetric initial brain seed. We then registered all the brains to the seed brain using the CMTK registration toolbox ^19, 20^. We used linear, followed by non-linear registration. We then generated a new average intensity and average deformation seed brain using an active deformation model (implemented in CMTK as avg_adm function). We iterated this process, obtaining at the end of the 5th iteration the *in vivo* intersex atlas (IVIA).

### Construction of IVIA-IBNWB bridging registrations

From all the fix-brain atlases with anatomical labels (IBNWB and JFRC), we chose the IBNWB atlas based on the similar nsyb-GFP signal coverage (neuropil, fiber bundles, and cell body rind) compared to the IVIA atlas (pan-neuronal mCD8-tdTomato). Despite this similarity, the two atlases have dissimilar distributions of fluorescence intensities per anatomical region and differences in signal to noise ratio (much higher for IBNWB), limiting the accuracy of intensity-based registration algorithms. Therefore, we used the rigid and non-rigid point set registration algorithm Coherent Point Drift (CPD) ^21^. We chose the surface of the antennal lobe (AL), antennal mechanosensory and motor center commissure (AMMCC), anterior optic tract (AOT), great commissure (GC), wedge commissure (WEDC), mushroom body (MB, includes mushroom body lobes, peduncle and calyx), posterior optic commissure (POC), protocerebral bridge (PB), lateral antennal lobe tract (lALT), posterior cerebro-cervical fascicle (pCCF), superior saddle commissure (sSADC), and the whole central brain as source of points to compare between atlases [Figure S3A]. We converted segmented binary images from each of these brain regions to triangular surface meshes via isocontour thresholding at isovalues of 0.9, generating a set of vertices and edges per brain region to be used for registration. We applied a rigid (translation, rotation and scaling) followed by a non-rigid CPD registration in both directions (IVIA-to-IBNWB, and IBNWB-to-IVIA) between the two sets of vertices across all brain regions. For the non-rigid registration, we explored the space of the two main meta-parameters *beta* (gaussian smoothing filter size) and *lambda* (regularization weight). We chose the non-rigid transformation (for either direction) with the highest jaccard index across all segmented brain regions that had spatially smooth deformations (visually inspected) [Figure S3B-C].

Using the IVIA-to-IBNWB transformation, we mapped anatomical segmentation of neuropils and neurite bundles ^22^ to the IVIA coordinates [Figure 1G]. In this segmentation, the AMMC would also include JON projections. In addition, we merged the mushroom body accessory calyx to the mushroom body calyx, due to its small size relative to average ROI volume.

### Registration accuracy

i) IVIA within-atlas accuracy: we segmented stalks from pC1, pC2l, pC2m, and pM7 neurons (n = 20, 10, 12, and 7 hemibrains for pC1, pC2l, pC2m, and pM7 respectively), which were consistently identified across individuals in flies expressing GCaMP6s via Dsx-GAL4 or CsChrimson via NP2631-GAL4 and FruFLP combo. These stalks covered different dorso-ventral regions from the posterior half of the brain [Figure S2B-i]. We measured the within-atlas registration accuracy as the standard deviation of each trace class from the mean trace (all traces across animals). The overall within-atlas accuracy was calculated as the standard deviation across all trace classes [Figure S2B-ii]. This estimate is the sum of biological variability and algorithm-associated error. Although we obtained cellular accuracy (∼4.3 μm), this is slightly lower compared to values reported for fixed brain atlases (2-3 µm) (Chiang et al., 2011; Jefferis et al., 2007; Peng et al., 2011; Yu et al., 2010), and this might be due to both a richer spatial pattern and higher signal-to-noise ratio of the BRP (nc82) antibody staining in fixed brain experiments.

ii) IVIA and IBNWB between-atlas accuracy: we used segmented pC1 stalks from IVIA (described above in i)) and compared them to pC1 neurons originally collected in the FCWB atlas (imaged via Fru-GAL4 expression of GFP ^23^), which were mapped to IBNWB (error of the transformation between FCWB and IBNWB is assumed to be negligible). Similar to i), we calculated the mean pC1 trace for each atlas (µ-pC1-IVIA and µ-pC1-IBNWB) and the within-atlas accuracy (σ-pC1-IVIA and σ-pC1-IBNWB) [Figure S3C]. We measured the IVIA-to-IBNWB registration accuracy as the difference between the σ-pC1-IBNWB and the standard deviation of pC1-IBNWB traces mapped to IVIA relative to µ-pC1-IVIA. On the other hand, we measured the IBNWB-to-IVIA registration accuracy as the difference between the σ-pC1-IVIA and the standard deviation of pC1-IVIA traces mapped to IBNWB relative to µ-pC1-IBNWB.

### Anatomical cataloging of central brain activity

For each fly, we combined all stimuli modulated ROIs and binarized these volumes, obtaining binary auditory voxel maps per fly. Auditory maps were then transformed to IVIA atlas coordinates. To account for the known fly-to-atlas error, we dilated these transformed maps ([1.25, 1.25, 2] µm^3^ in XYZ). We then summed these maps across flies, generating a density volume [Figures 2C, S1B, S4B-C, S5E, S7B, and S9A-B]. To spatially map auditory activity, we transformed neuropils and neurite bundle segmentation from IVIA coordinates to imaged volume coordinates. We assigned ROIs to a given neuropil or neurite bundle using the following criteria: i) ROIs must have at least a 15% volume overlap with its assigned neuropil or neurite bundle, ii) if overlapping with multiple neuropils or neurite bundles, ROIs are assigned to the neuropil or neurite bundle with the highest overlap. For auditory ROIs, we then quantified the volume they represent relative to the neuropil or neurite bundle for each fly (we used ROI, neuropil, and neurite bundle volume in IVIA space) [Figure 2E and S4E].

### Functional clustering of auditory responses to pulse, sine and white noise stimuli

For all auditory modulated ROIs from intact and *iav*^1^ flies (19,389 ROIs from 21 male (4 male flies carried *iav*^1^) and 17 female flies expressing tdTomato and GCaMP pen-neuronally) presented with the coarse tuning protocol (this includes pulse, sine, white noise, natural song, and P_fast_-like pulse, see Acoustic stimuli), we calculated the median response to each stimulus across trials, including 10 seconds before stimulus onset to 10 seconds after stimulus offset. We concatenated the median signal across pulse, sine, and white noise stimuli only (183 time points) for each ROI and z-scored it. We then hierarchically clustered ROI’s signals (pooled from all intact and *iav*^1^ flies) using a Euclidean distance metric and inner square distance metric between clusters (Ward’s method). This clustering split ROI responses into inhibitory and excitatory responses (first branching of the hierarchical tree), we then chose a distance (distance of 136) that would split both inhibitory and excitatory responses into the smallest number of distinct responses types (18 response types). We also required that each cluster be present in at least 16 flies (half the whole data set). Although we could further split these clusters, we stopped at 18 given that sub-clusters within each of the 18 clusters were very similar to the parent cluster, suggesting over-segmentation [Figure 3A].

### Response type kinetics and tuning

For each mean trace per response type, we measured adaptation, half-time to peak, and decay of auditory responses [Figure 3C-E]. Adaptation was measured as the percentage of absolute baseline subtracted signal 2 seconds before stimulus offset with respect to signal 2 seconds after stimulus onset (‘adaptation index’). Therefore, adaptation indices below 100% indicates some degree of adaptation (the lower the adaptation index, the higher the adaptation), while adaptation indices above 100% indicates a sustained increase in response (the higher the adaptation index, the higher the increase). Half-time to peak was measured as half the time to reach the peak signal during stimuli presentation. Decay time constant of auditory responses was determined by fitting an exponential to the 20 values after stimulus offset (10 seconds), except for response types 2 and 3, for which we fit an exponential to the 20 values after positive rebound.

To determine the coarse tuning of response types to pulse, sine or white noise, we measured the magnitude of stimulus-evoked responses. Response magnitude was defined as the absolute 80^th^ or 20^th^ percentile with the greater absolute value of baseline subtracted signal from stimulus onset to 2 seconds after stimulus offset (baseline was defined as the mean signal from -4 to -0.5 seconds relative to stimulus onset, dividing the response magnitude by the standard deviation during baseline gave us the signal-to-noise ratio). Response magnitudes were normalized by the maximum value across stimuli and multiplied by 100 to have units in percentage (%). We considered a response type as tuned to a particular stimulus if the response magnitude to the other stimuli was below 85% of the response to the preferred stimulus [Figure 3F-i-iii], otherwise it was considered non-selective [Figure 3F-iv].

### Clustering of tuning curves

We generated tuning curves to the ‘feature tuning’ stimulus set [Figure 4]. For these experiments, we sampled a fraction of each neuropil selected (24.4, 46.2, 10.5, 83, 39.8, 99.3, 63.6, and 91.8 % for the AMMC, SAD, GNG, WED, AVLP, PVLP, PLP and LH, respectively). For all auditory modulated ROIs from intact flies (10,970 ROIs from 10 male and 11 female flies) presented with the feature tuning protocol, we calculated the median response to each stimulus across trials, including 4 seconds before stimulus onset to 4 seconds after stimulus offset, and z-scored it. We measured the magnitude of responses to auditory stimulus for each ROI. Baseline was defined as -2 to -0.25 seconds from stimuli onset, we then calculated the mean µb of the signal during this time. Response magnitude was defined as the absolute 80^th^ or 20^th^ percentile with the greater absolute value of µb subtracted signal from stimulus onset to 2 seconds after stimulus offset. For each ROI, we generated a tuning curve by concatenating absolute response magnitude across presented sine song at different carrier frequencies, and pulse song at different carrier frequencies, pulse pause and pulse durations. We then hierarchically clustered ROI’s tuning curves using a Euclidean distance metric and inner square distance metric between clusters (Ward’s method). Examination of the tree revealed that tuning curves divide into three main classes, tuning to sine, tuning to pulses, and non-selective. We then reduced the distance threshold (distance of 60) until we split these three tuning types into the smallest number of visually distinct clusters [Figure 4C].

### Measuring activity diversity across brain regions

We collected ROIs belonging to the same neuropil across flies (separately for intact or *iav*^1^ flies). These ROIs, were sorted by their response type within a neuropil. Using the volume of each ROI, we calculated the total number of voxels for each response per neuropil, obtaining a distribution of voxel count per response type for each neuropil (we use ROI volume in IVIA space). These distributions were then max-normalized within neuropils; these were the values shown in Figure 3B. We then calculated the entropy for each response type distribution as a measure of activity diversity, regions with sparse response types will have a lower entropy value than regions with more broad or uniform distribution.

### Measuring trial-to-trial variability

To evaluate the variability of auditory responses across trials, we measured the residual of the ROI activity variance explained by the mean ROI activity across trials for pulse, sine and white noise stimuli (‘variability index’). Each trial included 10 seconds before stimulus onset to 10 seconds after stimulus offset. We concatenated the signal across stimuli and trials (183 time points) for each ROI and z-scored it. We computed the variability index for each ROI and then sorted these indices by neuropil [Figure 5B]. We did an all-to-all comparison of trial variability across neuropils using two-sample t-tests (we only included neuropils with at least 4 flies). We determined the significance of each comparison by bootstrapping. To correct for multiple comparisons, p-values were corrected using the Benjamini-Hochberg (B-H) FDR correction algorithm (FDR = 0.01).

### Measuring across-individual variability

We calculated response type distributions per neuropil (see Figure 3), but for each fly separately, obtaining an array of vectors of 18 dimensions per neuropil per fly. We normalized each vector to unit norm. We then use the cosine of the angle between normalized vectors as our measure of similarity of response distributions (‘similarity index’). Response distributions contained values greater than or equal to 0, restricting the distribution of similarity indices to the 0-1 range. We computed the similarity index between all possible combinations of fly pairs within a neuropil, generating a distribution of similarity indices for each neuropil. We compared these distributions across neuropils (for those with data from at least 4 flies), and significance of differences was determined using two-sample t-tests. We determined the significance of each comparison by bootstrapping. To correct for multiple comparisons, p-values were corrected using the Benjamini-Hochberg (B-H) FDR correction algorithm (FDR = 0.01). This correction assumes independence of positive dependency between tests.

### Measuring sex-specific differences in activity

Similar to the previous section, we calculated response type distributions per neuropil for each fly separately. These distributions were then sorted by sex. We then evaluated differences in probability of each of the 18 response types across sexes for each neuropil. We measured the effect size using the Cohen’s d [Figure S8C]. Within a neuropil, we determined the significance of female to male differences for each response type using two-sample t-tests [Figure S8B-C]. To correct for multiple comparisons within a neuropil, p-values were corrected using the Benjamini-Hochberg (B-H) FDR correction algorithm (FDR = 0.01).

### Identification of stimuli-modulated motor behaviors

Similar to the previous section (Identification of stimulus-modulated ROIs), we linearly modelled each motor variable (velocity and motion energy component, X(t)) as a convolution of the stimuli history and a set of three filters (one per stimulus). Filters were estimated using ridge regression ^16^, with a filter length of 10 seconds. Pearson correlation coefficient between raw motor variable (X(t)) to its linear prediction (g(t)), Pearson correlation coefficient between shuffled motor variable (sX(t)) to g(t), and corrected p-values were calculated as in section: Identification of stimulus-modulated ROIs.

### Linear modelling of neural activity from stimulus and behavior

To measure the fraction of neural variance that could be predicted from stimulus presentation or behavior of the fly, we constructed a linear model y(t) = μ(t) + Beta*X(t) + ε(t) [Figure 6G]. The raw signal for each ROI (y(t)) was modelled as the sum of the stimulus component (the trial average across auditory stimuli pulse, sine and white noise, μ(t)), behavior component (behavior predictors - velocities and top 500 - times a set of weights, Beta*X(t)), and noise (ε(t)). Behavior predictors were smoothed over time (Gaussian kernel size of 10 time points and gaussian kernel standard deviation of 3) and downsampled to match sampling rate of recorded activity (2 Hz). We computed the weights (Beta) of behavior predictors using ridge regression ^16^. To quantify the explanatory power of the full model or reduced models (stimulus or behavior component only), we computed the explained variance using 5 fold cross-validation.

### Analysis of hemibrain connectome

To find connections between early auditory neuropils - AMMC/SAD and WED - and downstream neuropils, we used the new ‘hemibrain’ connectome ^24^ [Figure 2G]. The ‘hemibrain’ project reconstructed 26,157 neurons and synaptic connections in an EM volume of ∼⅓ of the *Drosophila* central brain. This volume includes a large portion of the AMMC, SAD, and WED. We downloaded the connectivity matrix of all 26,157 neurons (https://www.janelia.org/project-team/flyem/hemibrain), and wrote custom code in python to find neuropils postsynaptic to a query neuropil, connected via one or two synapses. We considered a connection between two neurons to be significant if they had more than 5 synapses.

---

**Figure S1.**
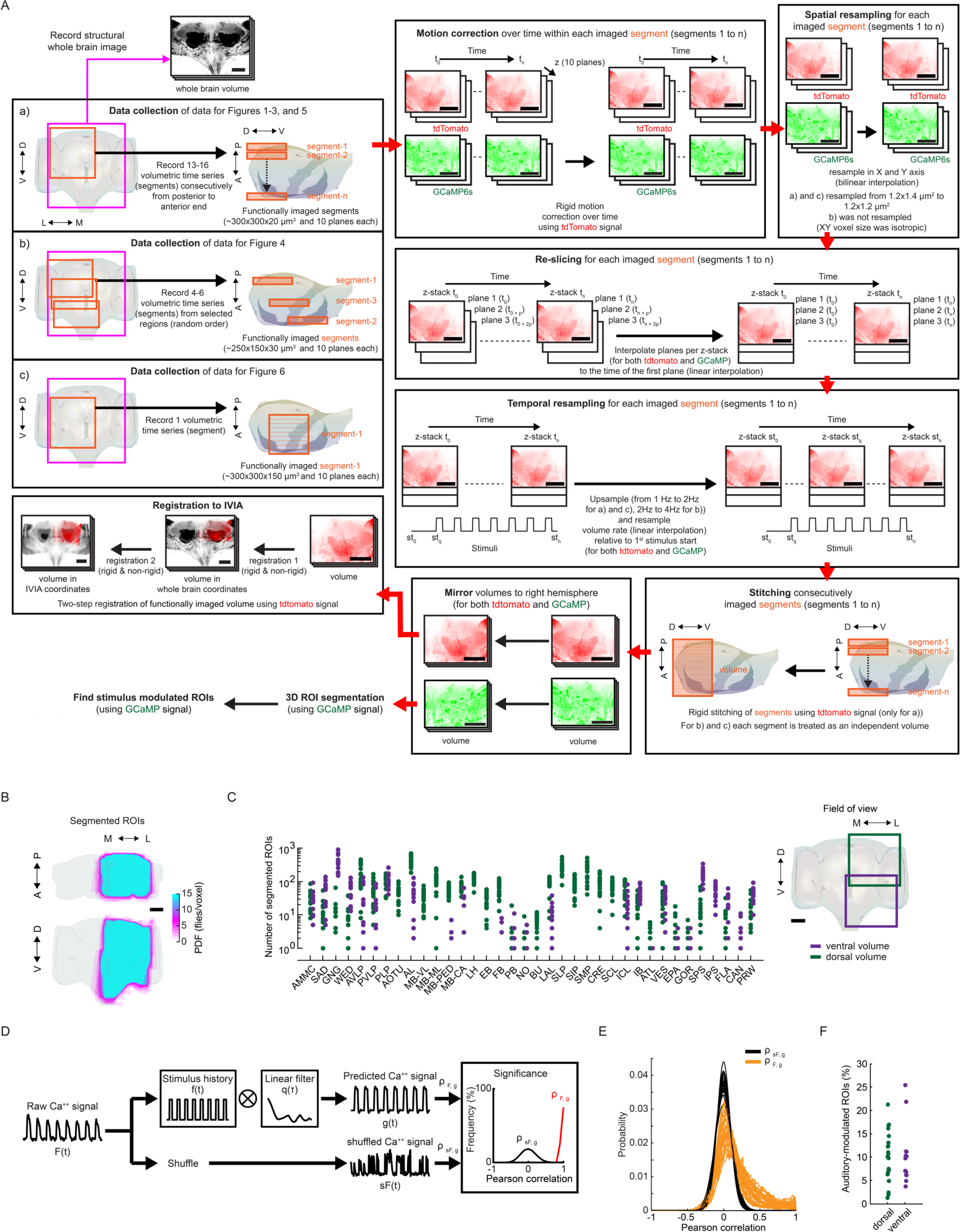
Spatial coverage of imaged activity and measurement of auditory-evoked activity. (A) Data collection and processing. We collected data using the following protocols: a) short volumetric recordings (∼10 min) of consecutive segments (from anterior to posterior axis), b) long volumetric recordings (∼30 min) of selected brain areas (non-consecutive), and c) short and coarse volumetric recording (∼15 min) of a single segment per fly that spans a large volume. For all these datasets we record a private whole brain structural volume (used later for registration). For all the data sets we recorded both tdTomato and GCaMP signal, and performed motion correction (using NoRMCorre) on each imaged segment using the tdTomato signal. These segments were then spatially resampled to have isotropic XY pixel size. This was followed by re-slicing of each Z-stack per segment (align time of all planes per Z-stack to the first plane imaged), and temporal resampling (aling Z-stack time relative to the start of the 1st stimulus and double the sampling rate). For protocol a), we stitched segments imaged consecutively (using NoRMCorre) obtaining a ‘volume’; for b) and c) each segment was treated as an independent volume. Volumes were mirrored to the right hemisphere, and the tdtomato signal was used for registration to the *in vivo* intersex atlas (IVIA). This is a two-step process, i) volumes per fly were registered to their own private whole brain volume (one per fly), and ii) whole brain volume was registered to the IVIA; registration i) and ii) were concatenated to map volumes to IVIA space. The GCaMP signal was used for ROI segmentation (via CaImAn), followed by identification of stimulus-modulated ROIs (see Figure S1D). (B) Maximum projection (in each dimension) of segmented ROIs from all imaged volumes (n = 33 flies, 185,395 ROIs) - ROIs from the left hemisphere are mirrored (see Figure S1A) such that all ROIs are projected onto right hemisphere. ROIs cover the entirety of the D-V and A-P axes. Color scale indicates the number of flies with an ROI in each voxel. (C) Number of ROIs across neuropils (see Figure 1F-G and Movie S1-4) sampled by ventral volumes or dorsal volumes (n = 33 flies, 185,395 ROIs). (D) Method for identifying stimulus-modulated ROIs. Raw Ca^++^ signal (F(t)) is convolved with the stimulus history (f(t)) and a set of filters per stimulus type (q(τ)) to generate the predicted Ca++ signal. Auditory modulation is measured by the cross-validated correlation scores (⍴_F,g_) between raw and predicted Ca^++^ signals. Correlation of shuffled Ca^++^ signal (sF(t)) to predicted signal (g(t)) is used to generate the null-distribution of correlation scores (⍴_sF,g_), which is used to determine significance. (E) Distribution of Pearson correlation coefficients of shuffled-vs-predicted signals (⍴_sF,g_) and raw-vs-predicted signal (⍴_F,g_) across all flies imaged (n = 33 flies). Unlike the distribution of ⍴_sF,g_, ⍴_F,g_ has a distribution with a long tail of positive correlation scores. ROis within the positive tail and outside the null distribution are considered to have significant stimuli modulation and selected as auditory ROis. (F) Frequency of auditory ROis separated by whether the ROI was segmented from a dorsal or ventral volume (n = 33 flies). Scale bars in all panels, 100 μm.

**Figure S2.**
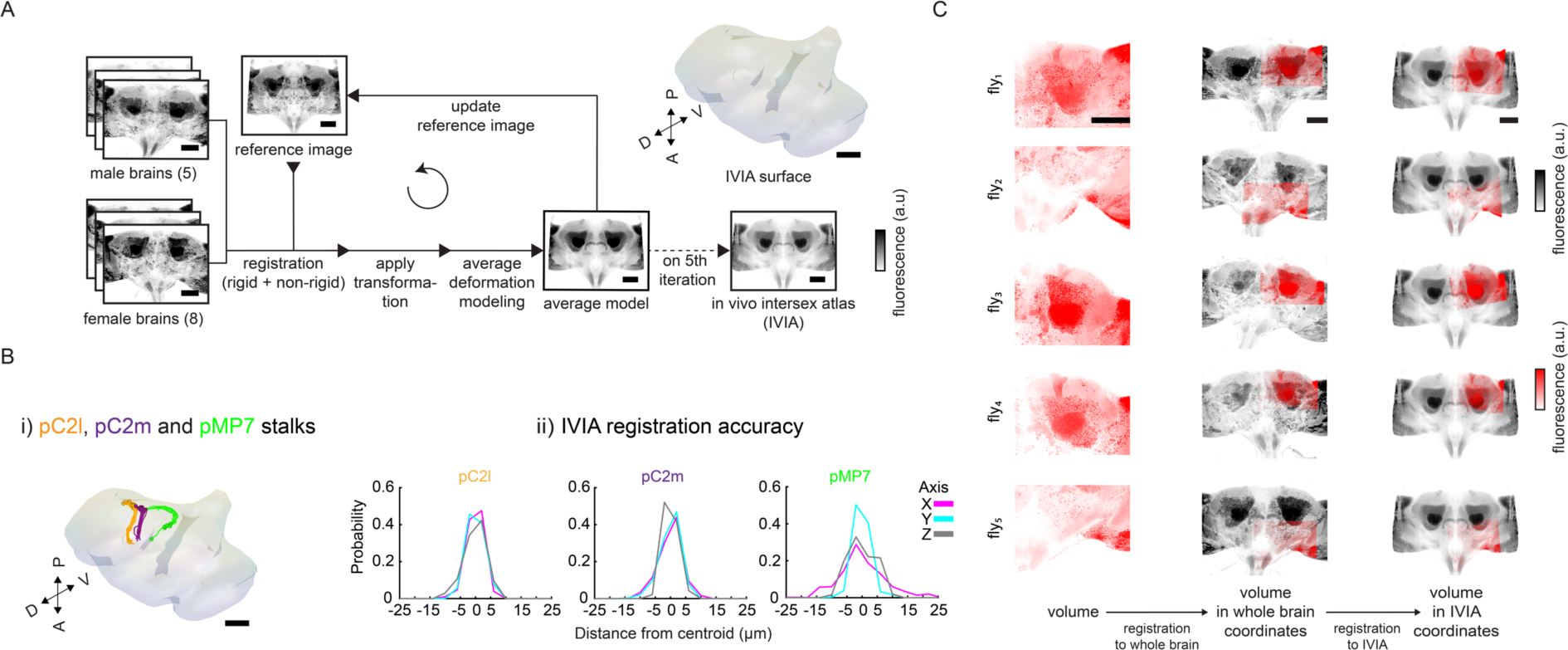
Building the *in vivo* intersex atlas (IVIA) (A) Generation of *in vivo* intersex atlas (IVIA): images of male (n = 5) and female (n = 8) brains expressing membranal tdTomato pan-neuronally are registered to a seed brain (reference image). Images are then transformed to generate an average model image, which, after five iterations, produces the *in vivo* intersex atlas (IVIA). (B) IVIA registration accuracy: i) 3D-rendering of traced tracts of Dsx+ neurons (pC2m, pC2l), and Fru+ neuron pMP7, ii) Per-axis jitter (X, Y, and Z) between matched traced tracts of pC2m, pC2l, and pMP7 across flies (n = 10, 12, and 7 brains, respectively). (C) Example imaged tdTomato volumes (red scale) from different flies registered to the IVIA (right-most column, gray scale). Middle column is the intermediate registration of volumes to their own private whole brain atlas (as in Figure S1A). Scale bars in all panels, 100 μm.

**Figure S3.**
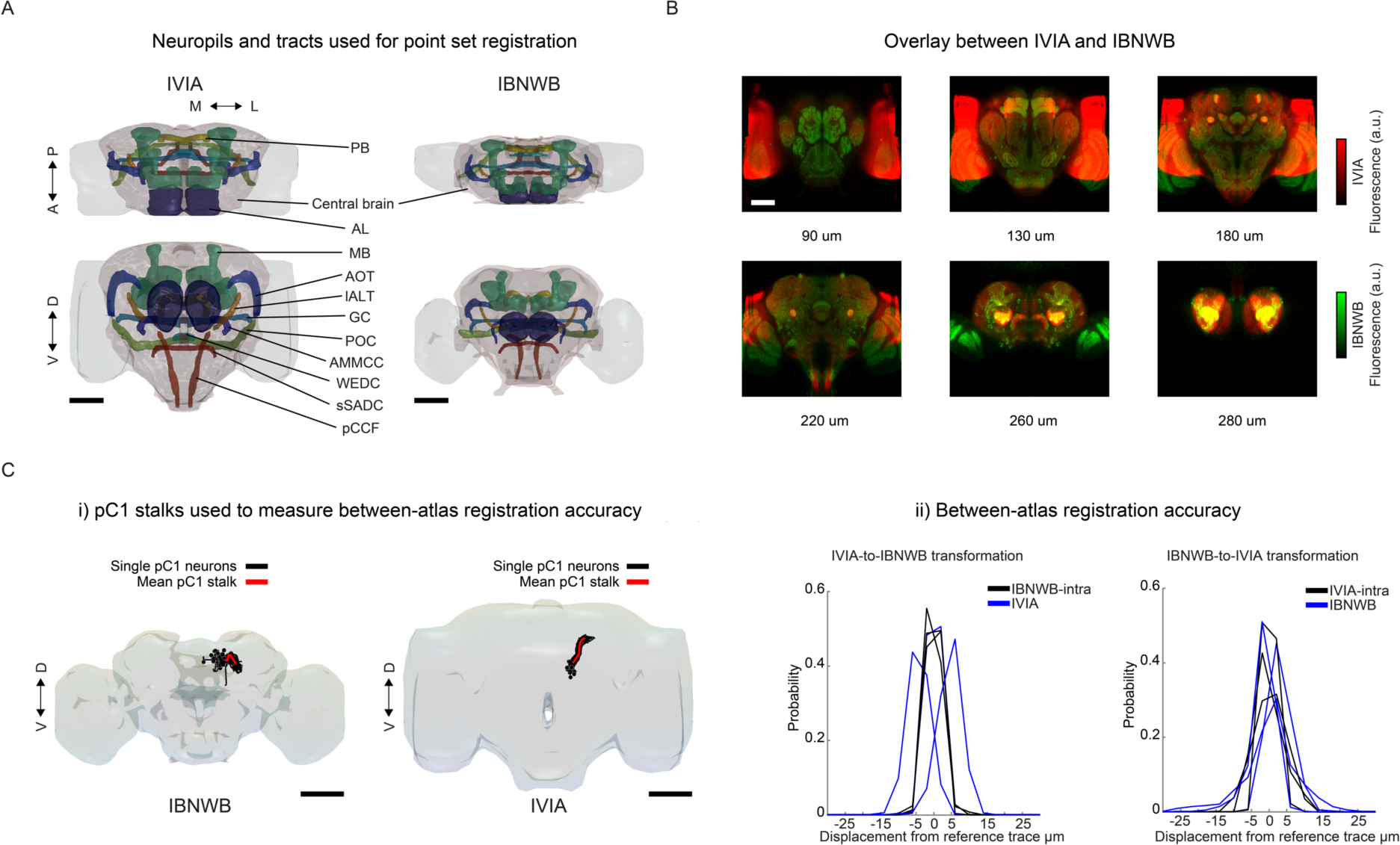
Registration between *in vivo* intersex atlas (IVIA) and fixed-brain atlas (IBNWB) (A) Central brain neuropils and tracts used for point set registration to morph the *in vivo* intersex atlas (IVIA) to IBNWB fixed brain atlas. Points from meshes of segmented antennal lobe (AL), mushroom body (MB) (includes mushroom body lobes, peduncle and calyx), protocerebral bridge (PB), antennal mechanosensory and motor center commissure (AMMCC), anterior optic tract (AOT), great commissure (GC), wedge commissure (WEDC), posterior optic commissure (POC), lateral antennal lobe tract (lALT), posterior cerebro-cervical fascicle (pCCF), superior saddle commissure (sSADC), and whole central brain were used to generate IVIA-to-IBNWB and IBNWB-to-IVIA transformations. (B) Overlay of IVIA (red) and registered IBNWB (in IVIA space, green) at different depths (90, 130, 180, 220, 260, and 280 µm). 0 μm is the most anterior section of the brain and 300 um the most posterior. (C) Atlas-to-atlas registration accuracy measured using pC1 stalks from IBNWB and IVIA. i) pC1 traces from IBNWB and IVIA atlases; black traces are single pC1 neurons (from IBNWB or IVIA, n = 70 and 20 pC1 traces respectively) and red trace is the mean reference pC1 stalk. ii) Between-atlas registration accuracy; IVIA- to-IBNWB transformation increases the jitter across all axes from the reference mean pC1 trace by ∼2.24 µm, while IBNWB-to-IVIA transformation increases the jitter by ∼2.8 μm. Scale bars in all panels, 100 μm.

**Figure S4.**
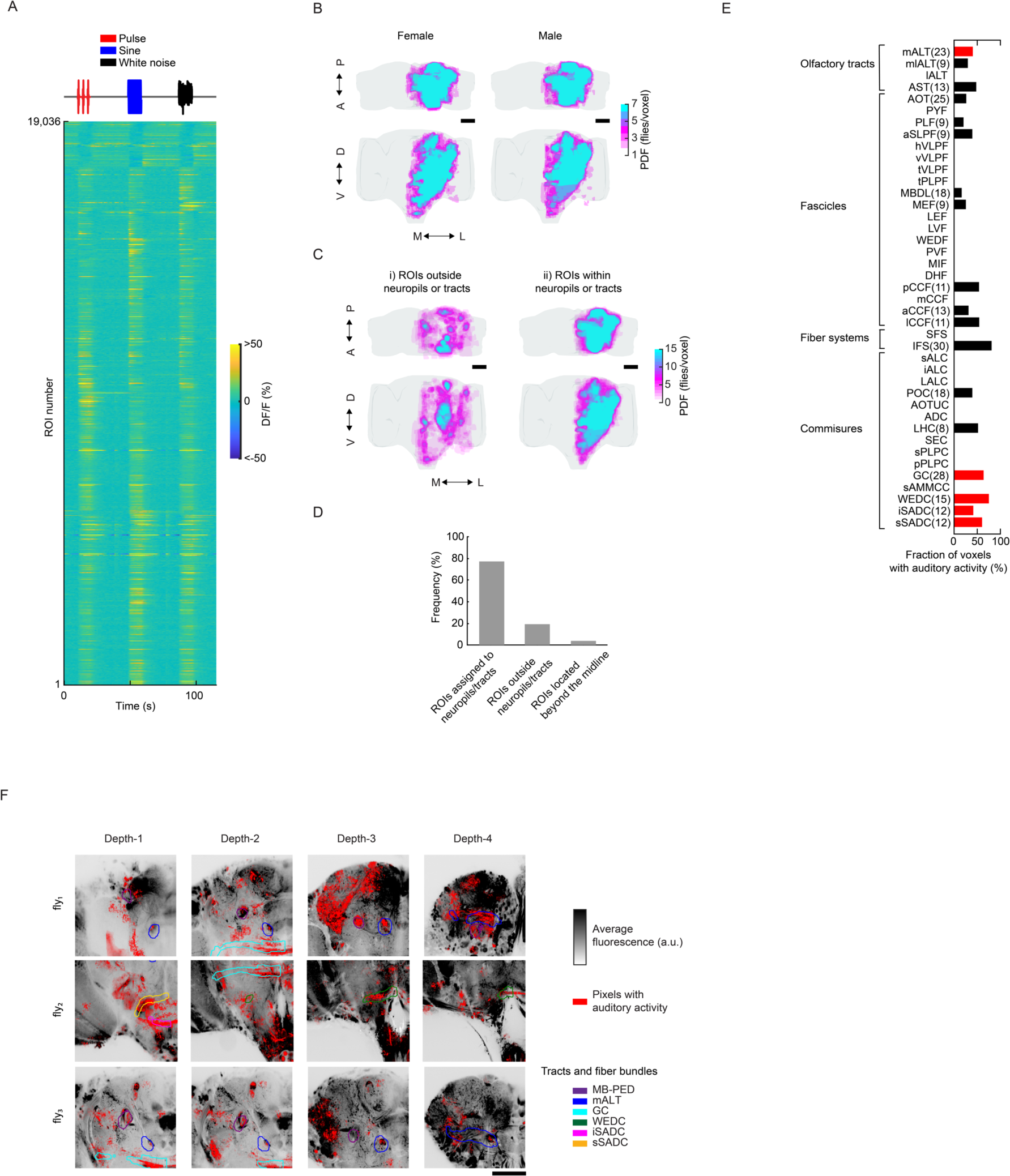
Auditory activity in neuropils and neurite tracts. (A) Median ROI responses (across 6 trials) to pulse, sine, and white noise stimuli (n = 33 flies, 19,036 ROIs) as in Figure 2A, but without z-scoring DF/F signal. (B) Spatial distribution of auditory activity across sexes. Maximum projections (from two orthogonal views) of the density of auditory ROIs in female (n = 17) or male flies (n = 16). (C) Maximum projection (from two orthogonal views) of the density of i) auditory ROIs outside neuropils or neurite tracts (n = 33 flies, 4,346 ROIs), and ii) auditory ROIs within neuropils or neurite tracts (n = 33 flies, 14,658 ROIs). Color scale for (B) and (C) is the number of flies with an auditory ROI per voxel. (D) Percentage of auditory ROIs within and outside neuropils and tracts, and beyond the midline. (E) Fraction of voxels with auditory activity by central brain neurite tracts; percentages averaged across 33 flies (a minimum of 4 flies with auditory activity in a given tract was required for inclusion). Red represents tracts that were clearly distinguishable from neuropil by visual inspection (see (F)). (F) For three flies, pixels with auditory activity (red) overlaid on time-averaged GCaMP6 fluorescence (grayscale). Several neurite tracts are indicated (planes from different depths are arbitrarily selected for each fly to highlight ROIs contained within neurite tracts). Scale bars in all panels, 100 μm.

**Figure S5.**
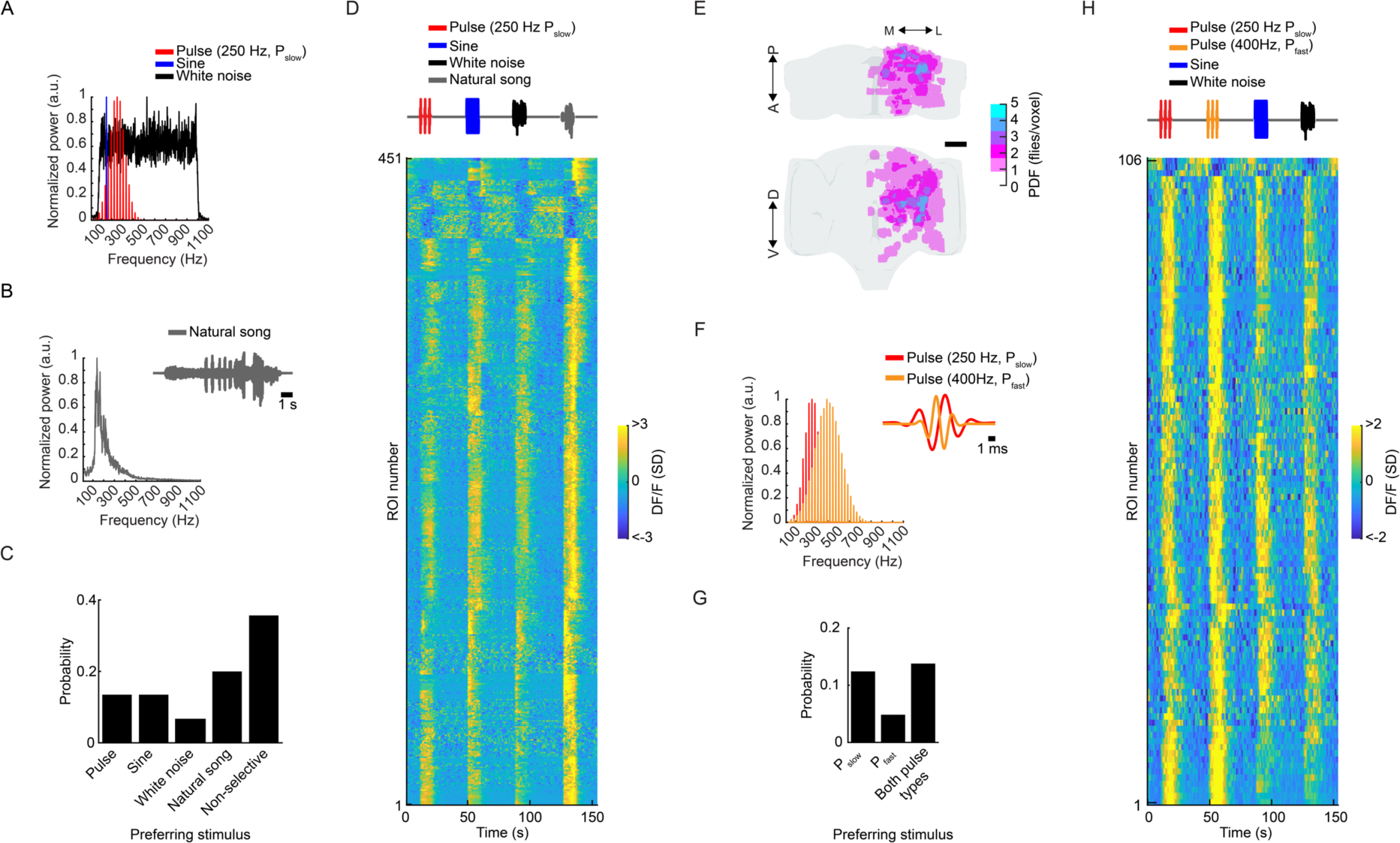
Auditory responses to additional song stimuli: natural song and fast pulses. (A) Spectral profile of auditory stimuli - pulse (P_slow_), sine, and white-noise - used to classify response types in Figure 3A, and their spectral features. (B) Spectral profile of natural song stimulus used. (C) Distribution of stimuli preference (to pulse, sine, white noise, and natural song) across auditory ROIs (n = 10 flies, 2,258 ROIs). Only ∼20% of auditory ROIs prefer natural song. Preference is defined by the stimulus that drives the maximum absolute response (at least 15% greater than the second highest response), as in Figure 3F. (D) Responses from auditory ROIs that prefer natural song (n = 451 ROIs). Each row is the median z-scored DF/F response across 6 across trials. Activity is plotted as the change in the standard deviation (SD) of the DF/F signal. (E) Spatial distribution of natural song preferring ROIs. Images are the maximum projection (from two orthogonal views) of the density of auditory ROIs with preference for natural song throughout the central brain (n = 10, 451 ROIs). Scale bars in all panels, 100 μm. (F) Spectral profile of P_fast_ and P_slow_ stimuli. (G) Distribution of stimuli preference to P_slow_, P_fast_, or broad preference for both pulse types across auditory ROIs (n = 2flies, 2,193 ROIs). Only ∼4% of auditory ROIs prefer P_fast_. Preference is defined as in (C). (H) Responses from auditory ROIs that prefer P_fast_ (n = 106 ROIs). P_fast_ preferring ROIs also show strong responses to P_slow_. Conventions same as in (D).

**Figure S6.**
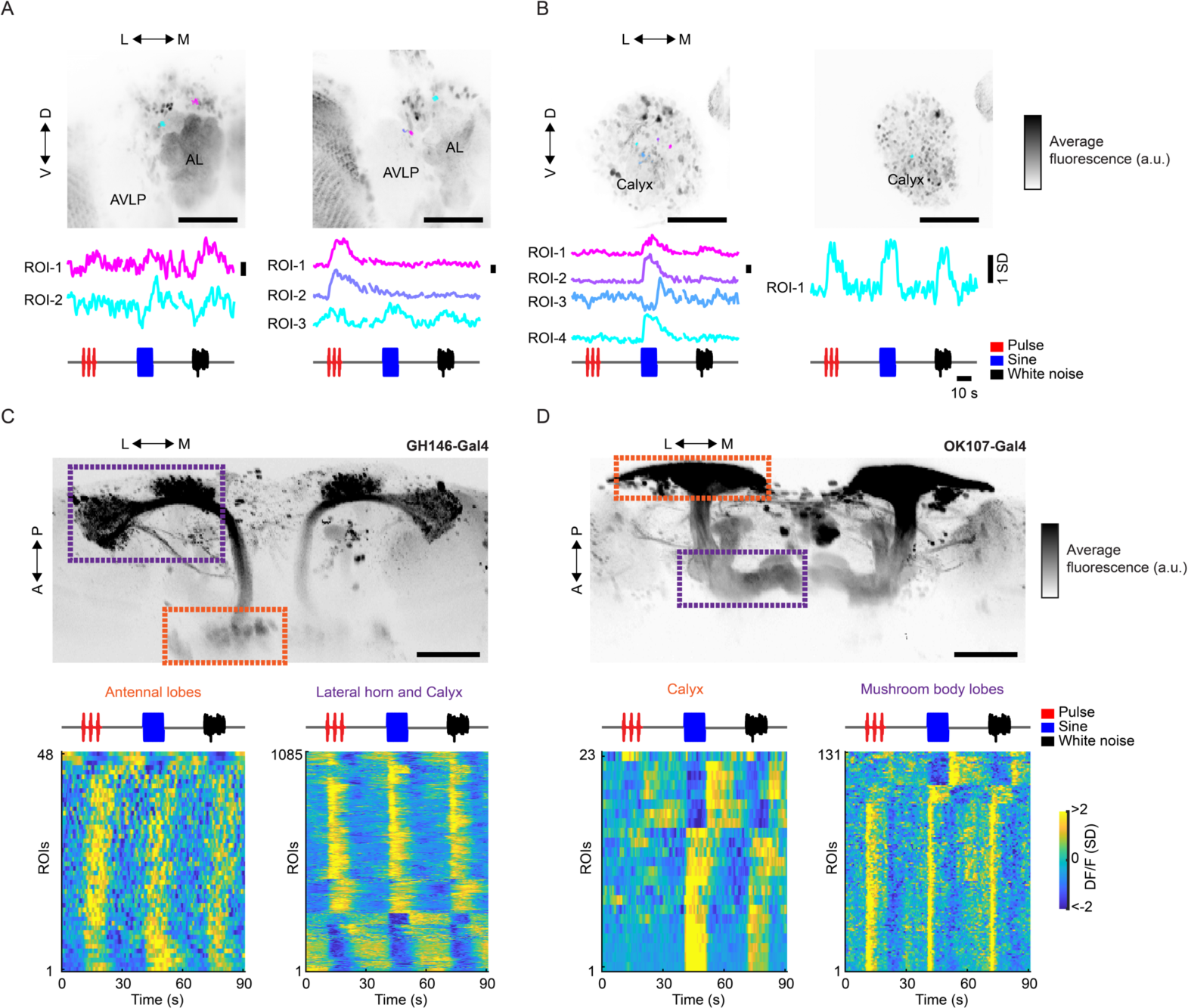
Auditory activity in olfactory neurons. (A-B) Auditory responses from (A) putative individual antennal lobe projection neurons (PNs), and (B) putative individual mushroom body Kenyon cells (KCs) from pan-neuronal recordings. Time-averaged GCaMP6s signal is shown in grayscale (over the entire experiment). Pixels belonging to individual ROIs and their corresponding time traces (median z-scored DF/F responses across 6 trials to pulse, sine, and white noise stimuli) are indicated in different colors. ROIs were drawn manually over the location of PN or KC somas. Activity scale bar unit is standard deviation (SD). (C-D) Auditory responses from (C) antennal lobe projection neurons, and (D) mushroom body Kenyon cells using cell type specific genetic lines (GH146-Gal4 and OK107-Gal4, respectively). Time-averaged GCaMP6s signal is shown in grayscale (over the entire experiment) - the compartments where activity was recorded from are indicated (data was collected and processed as we did for pan-neuronal data (protocol a) in Figure S1A) but the field of view was restricted to the region defined by the orange and purple boxes). Auditory ROIs are detected in all imaged compartments of antennal lobe projection neurons (n = 8 flies, 1,133 ROIs) and Kenyon cells (n = 4 flies, 154 ROIs). Each row is the median across 6 trials - all responses are z-scored and therefore plotted as the change in the standard deviation (SD) of the DF/F signal. Scale bars in all panels, 100 μm.

**Figure S7.**
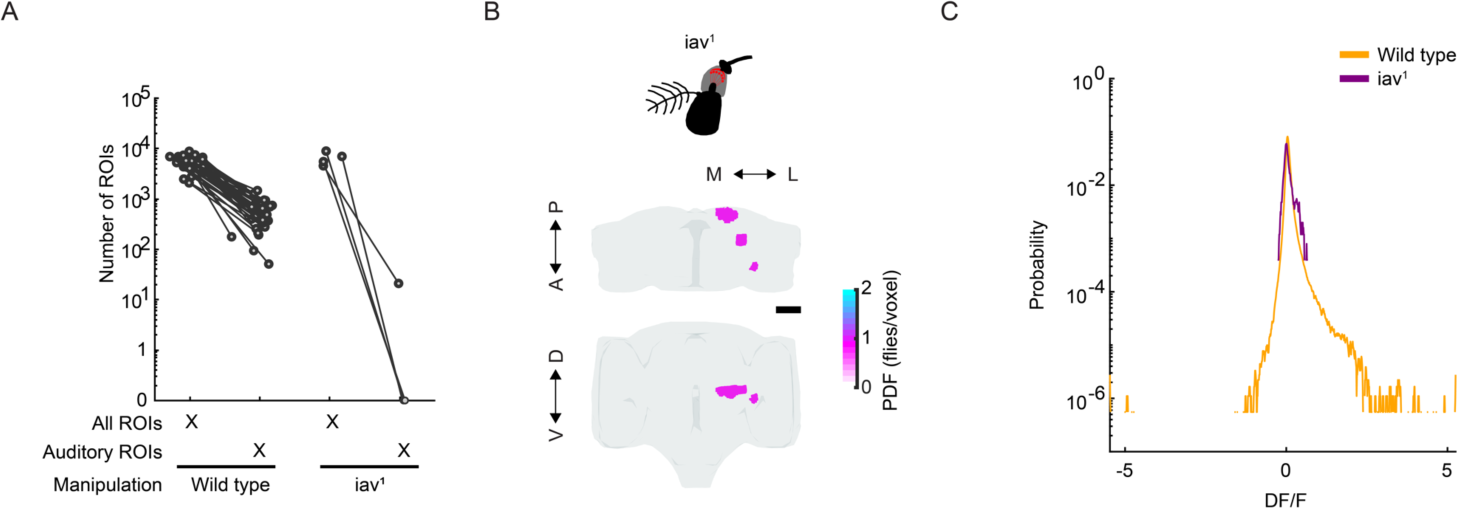
Brain-wide auditory activity is absent in deaf flies. (A) Number of detected auditory ROIs per fly in wild type (n = 33 flies) and *iav*^1^ flies (n = 4 flies). (B) Maximum projection (from two orthogonal views) of the density of auditory ROIs in *iav^1^* flies (all 21 ROIs come from 1 out of 4 flies imaged). These few ROIs were located in the mushroom body calyx (MB- CA), posterior VLP (PVLP), and the posterior lateral protocerebrum (PLP). Color scale is the number of flies with an auditory ROI per voxel. (C) Distribution of mean DF/F values (during stimuli) for auditory ROIs in wild type (n = 33 flies) and *iav^1^* flies (n = 4 flies). Scale bars in all panels, 100 μm.

**Figure S8.**
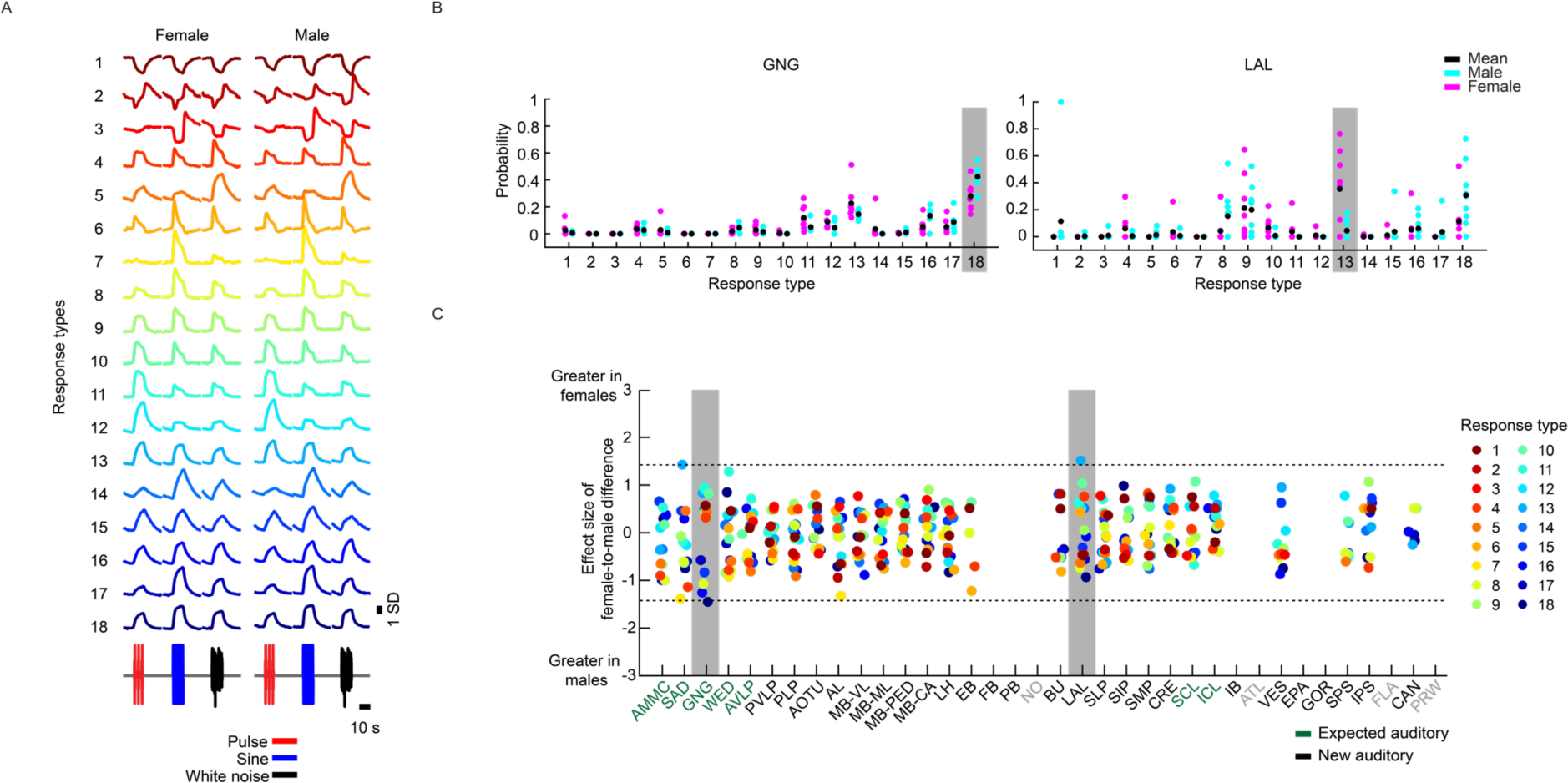
Auditory responses are not sexually dimorphic. (A) Hierarchical clustering of auditory responses into 18 distinct response types, and divided by sex. The mean response across ROIs belonging to each response type is shown. All responses are z-scored (as in Figure 3A), and therefore plotted as the standard deviation (SD) of the responses over time. (B) Probabilities of neuropil voxels with a given response type (separated by sex) in two neuropils, the gnathal ganglion (GNG) and the lateral accessory lobe (LAL). These are the only two neuropils with sex differences - males have a slightly higher probability of response type 18 activity in the GNG, while females have a slightly higher probability of response type 13 activity in the LAL. Each dot is the probability for one fly. (C) Sex-related differences in all response types across all neuropils. Each dot is the effect size (see Methods) of the difference in probability for each response type across sexes, color code is the same as (A). Neuropils with the greatest effect size are the GNG (response type 18) and LAL (response type 13), as highlighted in (B). However these differences are not significant (p > 0.05). Neuropils with no auditory activity are indicated in gray font.

**Figure S9.**
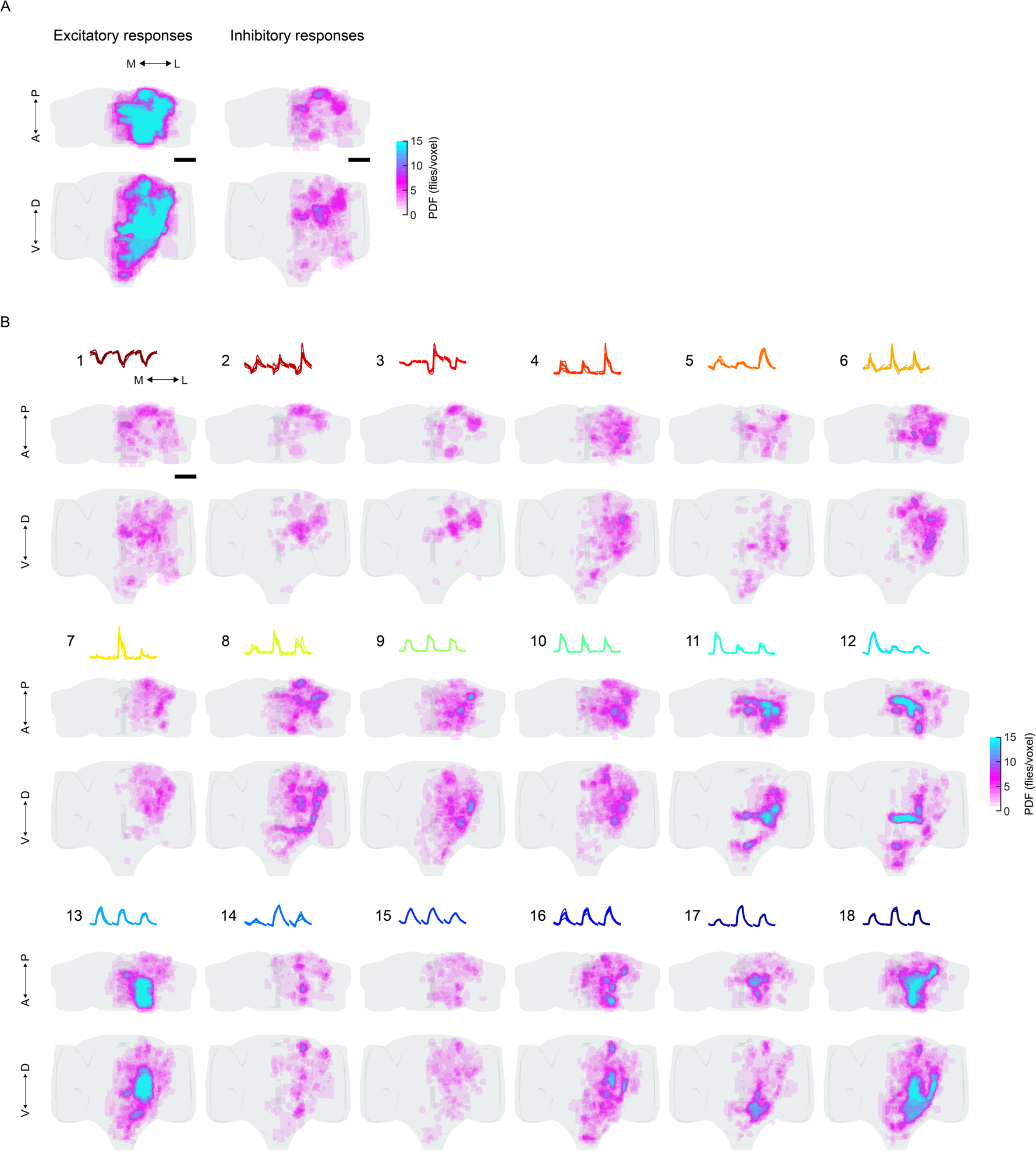
Auditory response types have distinct spatial distributions throughout the central brain. (A) Spatial distribution of excitatory vs inhibitory auditory ROI responses across the central brain. Images are the maximum projection (from two orthogonal views) of the density of excitatory and inhibitory responses across flies (n = 33). (B) Spatial distribution of auditory activity belonging to each response type (see Figure 3A) across the central brain. Images are the maximum projection (from two orthogonal views) of the density of responses across flies for each response type (n = 33). Color scale for (A) and (B) is the number of flies with an auditory ROI belonging to each category (excitatory, inhibitory, or response type) per voxel. Scale bars in all panels, 100 μm.

**Figure S10.**
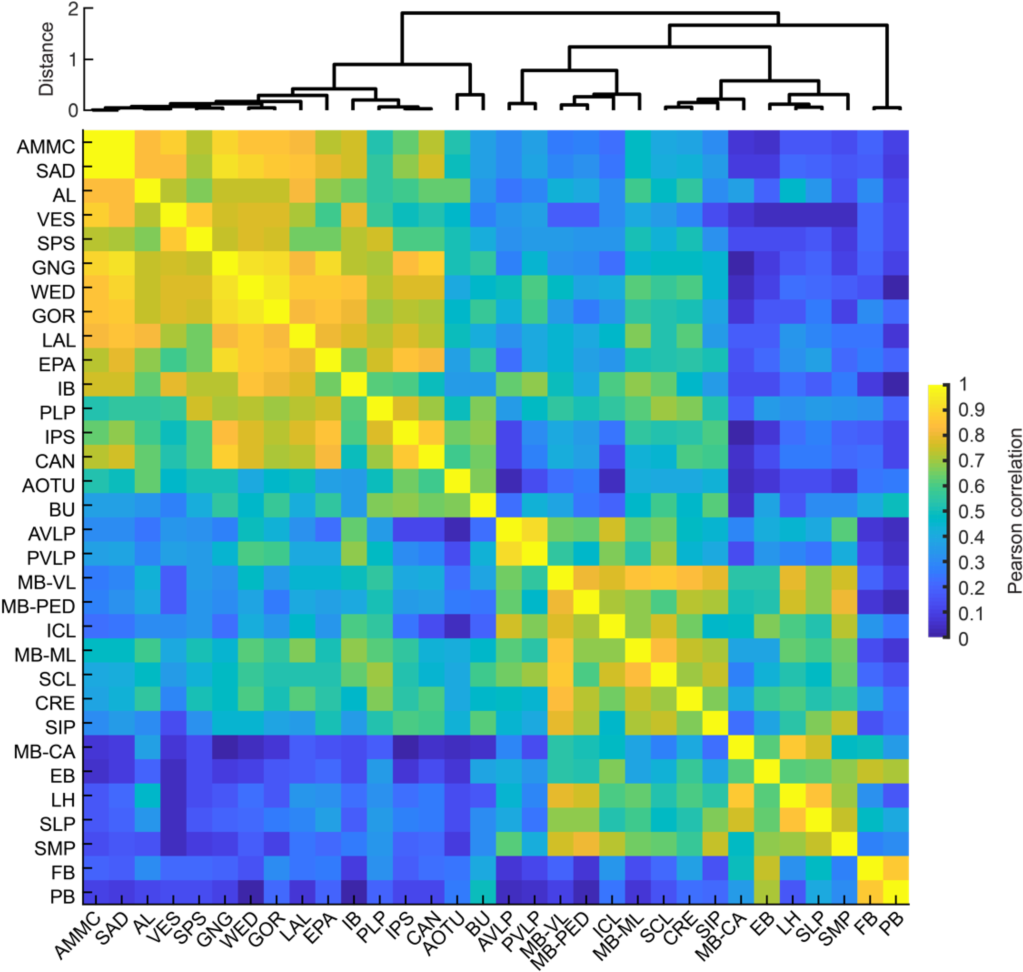
Similarity of auditory activity between neuropils. Pairwise Pearson’s correlation coefficients of response type distributions (as in Figure 3B) between neuropils. Neuropils are ordered based on the hierarchical clustering of the correlation matrix. Only positive correlation values are plotted for clarity.

**Figure S11.**
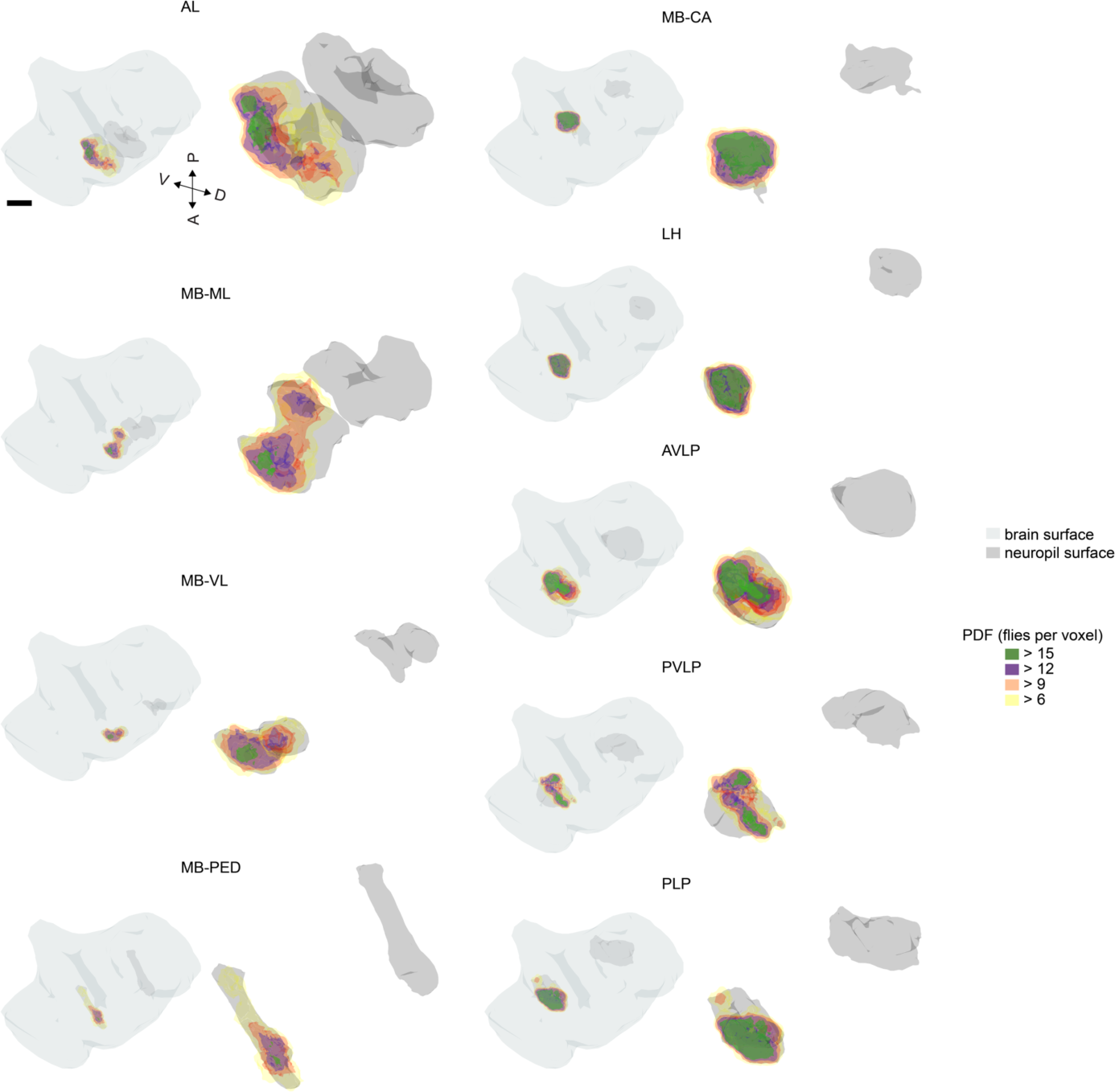
Auditory activity is compartmentalized within neuropils. Spatial distribution of auditory responses within selected olfactory (AL, MB-ML, MB-VL, MB-PED, MB-CA, and LH), visual (PVLP, and PLP) and mechanosensory neuropils (AVLP). Auditory activity is spatially restricted within each neuropil. Scale bars in all panels, 100 μm.

**Figure S12.**
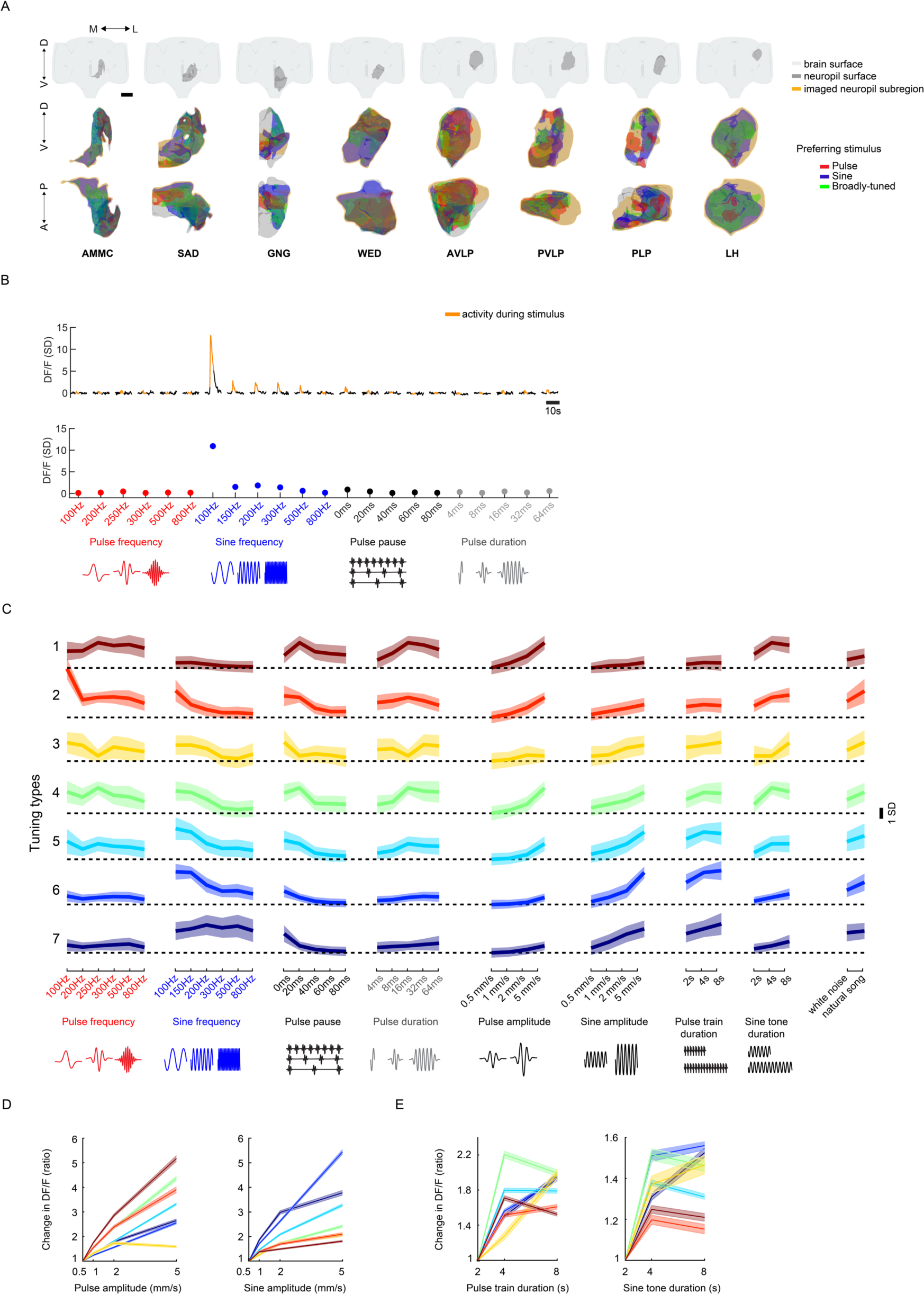
Related to Figure 4: Neuropil subregions imaged, definition of stimulus tuning, and tuning for additional song features. (A) Neuropil volume and subregion of neuropil imaged in Figure 4. Top row, location of the neuropil imaged (dark grey surface) relative to the central brain (light grey surface). Middle and bottom rows show maximum projections (from two orthogonal views) of imaged neuropil subvolume (orange surface) and the distribution of ROI stimulus selectivity (based on data in Figure 3B). Scale bar corresponds to 100 μm. (B) Example responses from one ROI to song feature stimuli (see Figure 6A). Top panel, Median (across trials) and z-scored responses to each song feature (responses are baseline subtracted, and baseline is defined as activity -4 to -0.25 seconds before stimulus onset). Orange timepoints correspond to activity during the stimulus. Bottom panel, response magnitudes (80^th^ or 20^th^ percentile of activity - for excitatory or inhibitory responses - during stimulus plus 2 seconds after) to each song feature calculated from the top trace (i.e. the tuning curve for this ROI). All responses are z-scored, so responses are plotted as standard deviation (SD) of DF/F value. (C) Tuning types as in Figure 6C, but plotting additional responses to pulses and sines of different amplitudes, varying pulse and sine train durations, and also to white noise and natural song (these additional responses were not used to cluster responses). Thick traces are the mean response magnitudes (calculated as in (B)) across all ROIs within each tuning type and shading is the standard deviation (n = 21 flies, 10,970 ROIs). Responses are plotted as standard deviation (SD) of DF/F value. (D) Auditory responses to different sine and pulse intensities, sorted by tuning type. Response magnitudes (calculated as in (B), but 80^th^ or 20^th^ percentile of the activity is measured during stimuli only) per ROI are normalized to the response to 0.5 mm/s stimuli. Thick traces are the mean normalized response magnitude, and shading is the s.e.m. (E) Auditory responses to different sine and pulse train durations. Response magnitudes (calculated as in (D)) per ROI are normalized to the response to 2 seconds stimuli. Conventions same as in (D).

**Figure S13.**
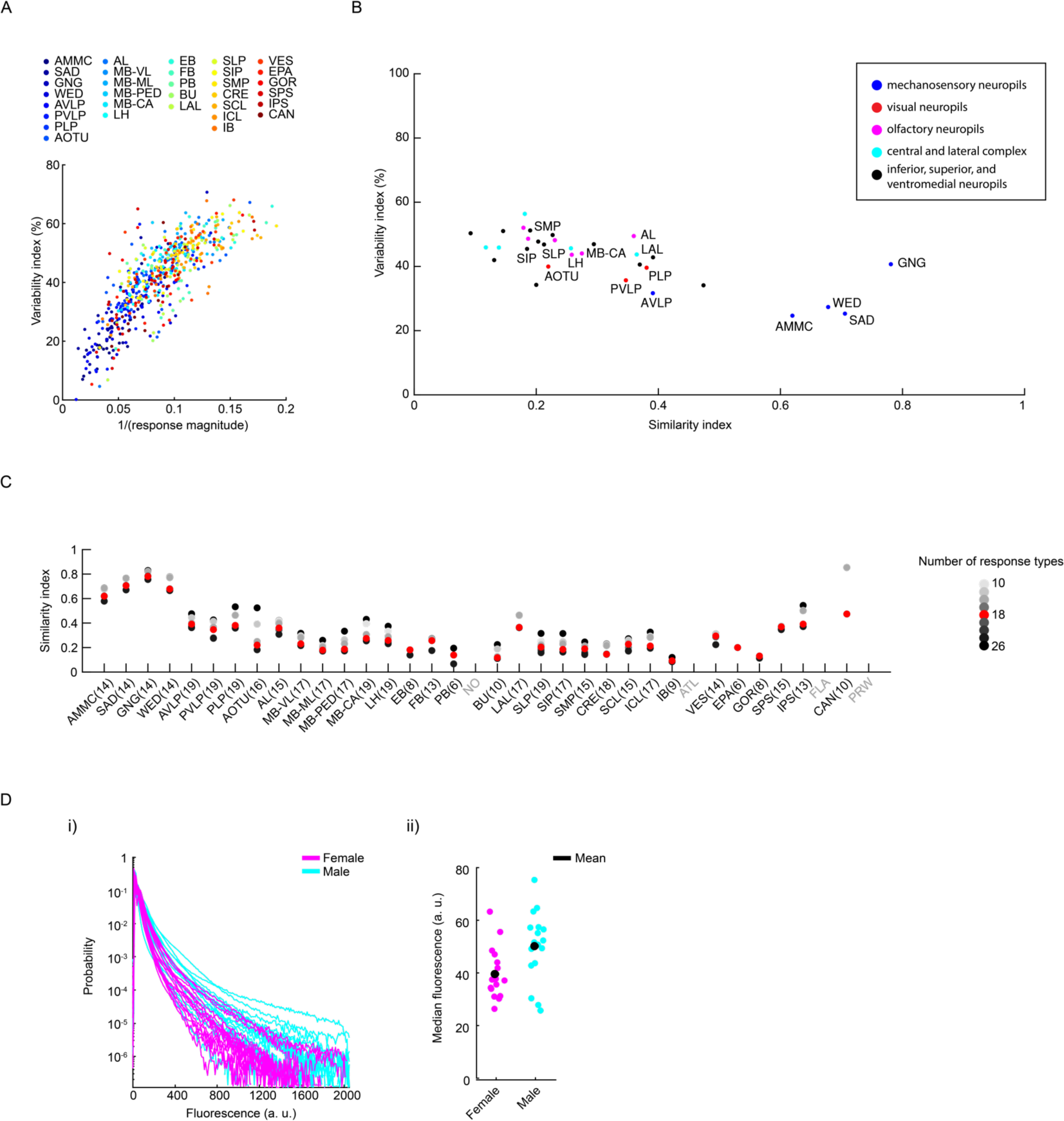
Across-trial and across-individual comparisons of auditory activity by neuropil. (A) Comparison of across-trial variability index (see Figure 5B) and response magnitude. Each dot is the mean variability index and response magnitude (80^th^ or 20^th^ percentile - for excitatory or inhibitory responses - of z-scored DF/F from stimuli onset until 5 seconds after stimulus offset) per neuropil for each fly (across all ROIs). Neuropils are color coded according to the legend. Variability index is inversely correlated with response magnitude. (B) Comparison of across-trial variability (see Figure 5B) and across-individual similarity (see Figure 5D). Each dot is the mean across-trial variability index and mean across-individual similarity index per neuropil. Groups of neuropils are color coded according to the legend. Early mechanosensory neuropils have low across-trial variability and high across-individual similarity. (C) Robustness of differences in similarity index across neuropils to the number clusters selected for hierarchical clustering of response types (see Figure 3A). Similarity index per neuropil (as in Figure 5D) is calculated for different numbers of clusters (from 10 to 26 - in Figure 3 we used 18 types (red)). (D) No systematic difference in distribution of time-average fluorescence (over the entire experiment) across individuals. i) Histograms of time-averaged fluorescence per individual, cyan and magenta correspond to male (n = 16) and female (n = 17) flies. ii) Median fluorescence per individual. Black dot corresponds to the mean (of median fluorescence) across flies.

**Figure S14.**
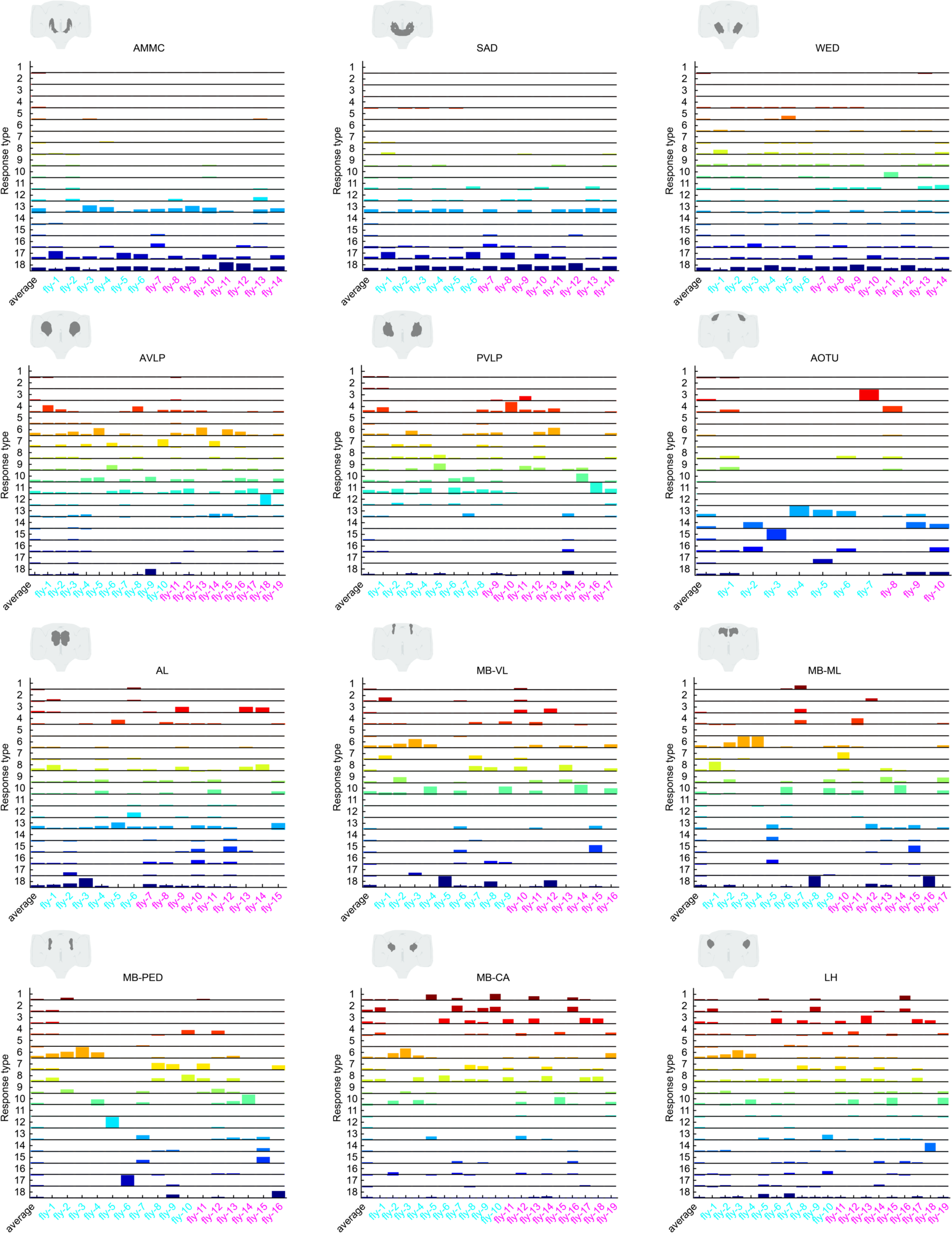
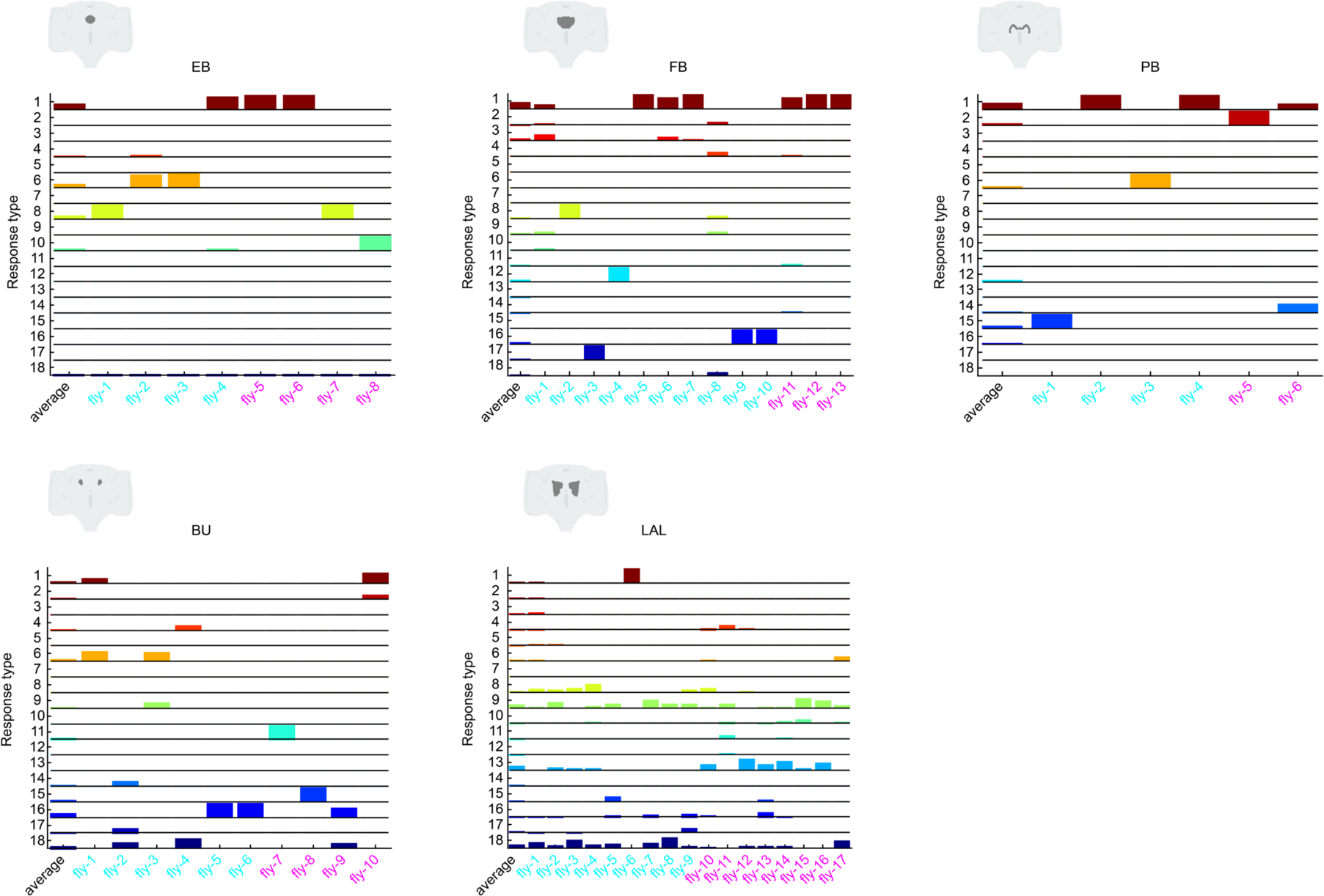
Distribution of 18 auditory response types per neuropil, separated by flies. Response type distributions, similar to Figure 5C, for all mechanosensory, visual, olfactory, central and lateral complex neuropils. For each neuropil, male and female flies are indicated in cyan or magenta, respectively.

**Figure S15.**
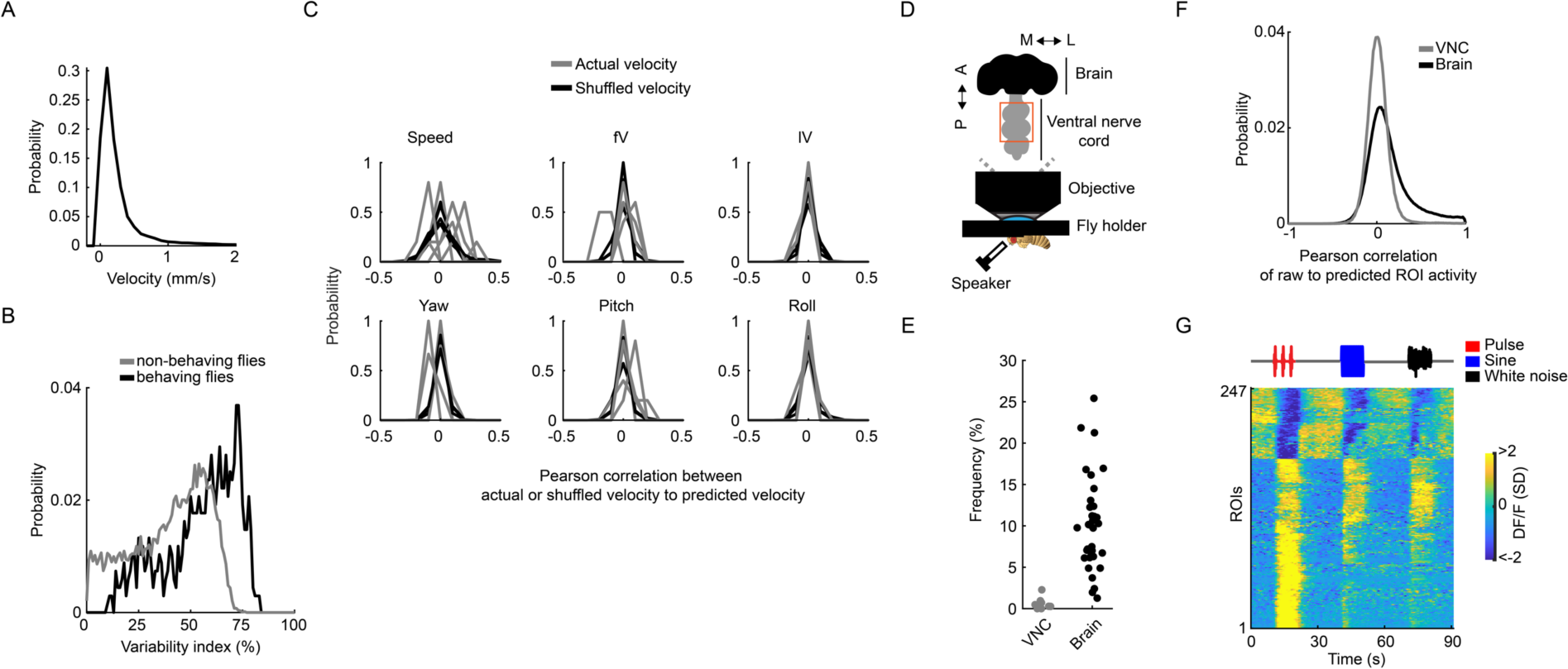
Behavior of head-fixed flies and minimal auditory responses in the *Drosophila* ventral nerve cord. (A) Distribution of speed across flies (n = 5 flies). (B) Distribution of across-trial variability of auditory ROIs in behaving (data used in Figure 6, n = 5 flies, 849 ROIs) and non-behaving flies (data used in Figure 5B, n = 33 flies, 19,036 ROIs). (C) Correlation between linearly predicted velocities (based on stimuli history) and actual or shuffled (see Methods) velocities. Each black or gray line is the probability per fly (n = 5 flies). (D) Schematic of VNC functional imaging during auditory stimulation. Similar to Figure 1A, but the dorsal side of the thorax is dissected to expose the dorsal side of the first and second segment of the VNC depicted inside the orange rectangle. (E) Frequency of auditory ROIs detected in the brain (n = 33 flies, 185,395 ROIs) and the VNC (n = 8 flies, 39,580 ROIs) using the same criteria as in Figure S1D. (F) Distribution of Pearson correlation coefficients of raw ROI activity to predicted ROI activity (based on stimuli history, as in Figure S1E) for ROIs recorded from the brain (n = 33 flies, 185,395 ROIs) and the VNC (n = 8 flies, 39,580 ROIs). (G) Responses of VNC auditory ROIs to pulse, sine, and white noise stimuli (n = 8 flies, 39,580 ROIs). Each row is the across-trial median and z-scored DF/F response to each stimulus (6 trials per stimulus). DF/F units are in standard deviation (SD).

## Movie legends

**Movies S1-S4** DF/Fo time-series movies (maximum intensity projected over entire z axis of the Drosophila brain) of male (S1-2) and female (S3-4) brain dorsal (S1 and S3) and ventral (S2 and S4) quadrants. Responses to pulse (250Hz), sine (150Hz) and white noise are shown. Pseudocolor scale is from 0 to 200 %. Movie speed is sped up 5X.

**Movie S5** Z-stack of the probability density of auditory ROIs (number of flies containing an auditory ROI at each voxel) throughout the central brain (n = 33 flies, 19,036 ROIs). Mechanosensory neuropils have grey contours, while the rest of the central brain neuropils imaged have red contours. Pseudocolor scale is from 0 to 15 flies/voxel.

## Abbreviation

AMMC: antennal mechanosensory and motor center
SAD: saddle
GNG: adult gnathal ganglion or SEZ
WED: wedge
AVLP: anterior ventrolateral protocerebrum
PVLP: posterior ventrolateral protocerebrum
PLP: posterior lateral protocerebrum
AOTU: anterior optic tubercle
AL: adult antennal lobe
MB-VL: vertical lobe of adult mushroom body
MB-ML: medial lobe of adult mushroom body
MB-PED: pedunculus of adult mushroom bod
MB-CA: calyx of adult mushroom body
LH: lateral horn
EB: ellipsoid body
FB: fan-shaped body
PB: protocerebral bridge
NO: nodulus
BU: bulb
LAL: lateral accessory lobe
SLP: superior lateral protocerebrum
SIP: superior intermediate protocerebrum
SMP: superior medial protocerebrum
CRE: crepine
SCL: superior clamp
ICL: inferior clamp
IB: inferior bridge
ATL: antler
VES: vest
EPA: epaulette
GOR: gorget
SPS: superior posterior slope
IPS: inferior posterior slope
FLA: flange
CAN: cantle
PRW: prow

## Notes

### Competing Interest Statement

The authors have declared no competing interest.

### Summary of Updates

Results and Discussion text, Figures, and Supplemental Figures updated. We added new data and analysis of neural recordings in behaving flies (Figure 6).

